# Longitudinal single-cell RNA sequencing of a neuroendocrine transdifferentiation model reveals transcriptional reprogramming in treatment-induced neuroendocrine prostate cancer

**DOI:** 10.1101/2025.04.25.650369

**Authors:** Funda Sar, Hee Chul Chung, Yen-Yi Lin, Dong Lin, Tunc Morova, Anne Haegert, Stanislav Volik, Robert Bell, Stephane LeBihan, Dogancan Ozturan, Hui Xie, Xin Dong, Rebecca Wu, Jing Li, Hsing-Jien Kung, Martin E. Gleave, Nathan A. Lack, Yuzhuo Wang, Colin C. Collins

## Abstract

Neuroendocrine transdifferentiation (NEtD) of prostate adenocarcinoma (PRAD) leads to aggressive neuroendocrine prostate cancer (NEPC). The LTL331 patient-derived xenograft (PDX) model consistently progresses to NEPC following castration, mimicking clinical responses to androgen-deprivation therapy. Here we tracked NEtD using longitudinal single-cell RNA sequencing (scRNA-seq) across eight time points in LTL331 from pre- to post-castration. Castration led to the loss of AR-high PRAD cells, expansion of AR-low populations, and emergence of an AR- and NE-negative (AR-/NE-) intermediate state that transitioned into NEPC. We delineate a model in which pre-EMT cells enriched in ciliogenesis and cell-adhesion pathways differentiate into EMT-like populations before branching into distinct ASCL1+ and ASCL1− NEPC states. The EMT-enriched intermediate, marked by progenitor and neural crest stem-cell genes, acts as a transition bridge, suggesting EMT-associated stemness underlies lineage plasticity. A terminal ASCL1-NEPC state also suggests that ASCL1 is not essential for NEPC maintenance. Gene regulatory analysis highlighted key regulators driving NEtD, including MSX1 and ASCL1 in early EMT-like and NEPC states, respectively. These transcriptional states were validated in both patient-derived bulk and scRNA-seq data. Our findings offer insights into intervention strategies to delay or prevent NEtD with the potential of identifying novel prognostic and therapeutic targets.

## Introduction

Globally, prostate cancer (PCa) was diagnosed in 1,466,680 men and killed 396,792 in 2022 (1). These numbers are expected to worsen rapidly in the coming years (2). Androgen deprivation therapy (ADT) that inhibits the androgen receptor (AR) signaling axis is the standard of care for most PCa patients. AR pathway inhibitors such as enzalutamide (3) extend the life of PCa patients. However, the onset of resistance in the form of castrate-resistant prostate cancer (CRPC) is universal, with a median survival time of 3-4 years (4,5). Resistance can result from reactivation of the AR signaling axis or through lineage plasticity where AR signaling is lost and prostatic adenocarcinoma (PRAD) cells give rise to a neuronal phenotype (6,7) characterized by neuronal markers such as chromogranin A. This process is called neuroendocrine transdifferentiation (NEtD). While the precise mechanisms of NEtD remain to be defined, lineage plasticity has emerged as an important mechanism of treatment resistance (see (8) for review). Treatment-induced NEtD is associated with profound changes in cell phenotypes ranging from a loss of luminal features to induction of neuronal phenotypes. There are significant gaps in our understanding of the mechanisms underlying NEtD (8). Neuroendocrine prostate cancers (NEPC) are now seen in up to 25% of patients with CRPC (9,10). There are no therapeutic options for NEPC, and it is rapidly fatal making identification of therapeutic targets vital. Prior efforts to understand NEtD using single-cell RNA-seq (scRNA-seq) profiled individual patients at a single time point (11–13). The lack of a time course coupled with tumor and patient heterogeneity-imposed challenges for elucidating the NEtD transcriptional continuum for these studies. Therefore, to gain deeper insight into the mechanism of NEtD and to identify potential therapeutic targets, we combined scRNA-seq with a unique patient-derived xenograft (PDX) NEtD model (14) in a longitudinal study.

The LTL331 model phenocopies the donor patient’s clinical course by undergoing NEtD (15) in response to host castration. Exome sequencing and whole transcriptome sequencing revealed it to be TP53, RB1, and PTEN negative (15) consistent with an NEPC phenotype (6). Thus, it is uniquely valuable because it allows tracking NEtD longitudinally from PRAD to NE enabling detailed dissection of the lineage plasticity phenomenon and identification of biomarkers and therapeutic targets. We used bulk genomics and transcriptomics (15,16) and proteomic (17) methods to characterize the NEtD process in LTL331 model. This resulted in the identification of mediators of NEtD including HP1α (18) BHC80 (19), BRN2/POU3F2 (20), PEG10 (15), ONECUT2 (21), SRRM4 (22), and H19 (16) all of which were validated in clinical cohorts underscoring the clinical relevance of this model. The establishment and maintenance of cell type-specific gene expression programs are regulated by lineage-specific transcription factors (TFs). AR and other related transcription factors (TFs) (e.g., ERG, FOXA1) are considered key drivers of PRAD development. In contrast, the loss of these TFs occurs concomitantly with the expression and activation of NE lineage TFs (e.g., BRN2/POU3F2, FOXA2, ASCL1) in terminal NEPC. To date, only a small number of NEPC-related TFs have been identified, and they are mainly reported in terminal differentiation states. The hypothesized driver TFs of NEtD remain largely unknown. Identification of these TFs will have profound implications for understanding the mechanism(s) of NEPC development and lead to more effective disease management.

To elucidate early events and regulatory mechanisms involved in NEtD, we conducted a longitudinal scRNA-seq analysis using the LTL331 model (15–17). The scRNA-seq data revealed a complex and dynamic response to castration, suggesting intricate regulatory processes underpinning NEtD. Specifically, castration induces expansion of small pre-existing cell populations at different stages of NEtD, characterized by low expression levels of AR and prostate-specific antigen (KLK3), accompanied by a concurrent decline in the dominant population that expresses high levels of AR and KLK3. We identified a pair of phenotypically distinct cell clusters that converge on a critical transition state marked by epithelial-mesenchymal transition (EMT), implying significant changes in cellular morphology and function during NEtD. From this EMT-like state, our analysis delineated bifurcating lineages, leading to the formation of early NEPC cell clusters followed by late and heterogeneous NEPC clusters. This finding suggests multiple adaptive pathways leading to and resulting in distinct NEPC subtypes, each potentially driven by different oncogenic and survival mechanisms.

To assess the clinical relevance of these heterogeneous clusters, we cross-referenced our findings with existing bulk and single-cell NEPC transcriptome studies. Our integrative approach allowed us to detect clusters similar to many of these NEPC subtypes in LTL331. Moreover, we identified novel transcription factors to understand their potential roles in NEtD, that may underly the transcriptional heterogeneity and lineage plasticity (21). This confluence of LTL331 insights with historical and clinical data underscores the complex biological dynamics of NEPC and offers a refined view of its molecular underpinnings, potentially guiding toward more effective therapeutic strategies.

## Materials and methods

### LTL331 PDX model

The LTL331 PDXs were obtained by grafting under the kidney capsules of non-obese diabetic (NOD) severe combined immunodeficiency (SCID) mice as previously described in (14). Tumor tissues were extracted at individual time points as described in (14,15) . Ethical guidelines in the Declaration of Helsinki were followed. Informed written consent was obtained from the patient using a protocol (#H09-01628), approved by the Institutional Review Board of the University of British Columbia (UBC). The animal studies protocols (#A17-0165) were approved by UBC’s Animal Care and Use Committee.

### Histopathology and immunohistochemistry

Preparation of paraffin-embedded tissue sections and immunohistochemistry were carried out as previously described in (23). Routine hematoxylin and eosin (H&E) staining was performed. For immunohistochemistry, a rabbit polyclonal anti-AR antibody (Affinity BioReagents), a rabbit polyclonal anti-PSA antibody (Dako), a mouse monoclonal anti-NCAM1 antibody (Cell Marque) were used.

### Tumor dissociation and single cell RNA sequencing

Each tumor was diced into small pieces using a scalpel (Bard-Parker) and then dissociated into a suspension of viable cells using human tumor dissociation kit (Miltenyi Biotec) according to manufacturer’s protocol. Following dissociation cells were resuspended in 1X PBS (Gibco) supplemented with 0.04% BSA (Milteyni Biotec). The cells were counted, and viability was assessed using 0.4% (w/v) Trypan Blue staining (Thermo Fisher). We proceeded to scRNA-seq with samples with a minimum of 80% viable cells. Cells were loaded onto the 10X Chromium controller to capture ∼10.000 cells. The single cell RNA libraries were prepared using Chromium Single Cell 3’ v3.1 reagents following manufacturer’s protocol and sequenced with an average read depth (47,760 reads/cell) using Complete Genomics DNB-Seq G400 platform (**Supplementary Table 1)**.

### Whole exome sequencing (WES) of LTL331 model

DNA extracted from four PDX tissue samples (preCx, 16wk, 20wk and 22wk post castration) was used to prepare whole exome sequencing (WES) libraries according to the KAPA HyperCap workflow 3.0 (Roche Diagnostics) but utilizing the MGIEasy DNA adapters for MGI sequencing. For pre-capture multiplexing, the amplified libraries were multiplexed to a combined mass of 1.5 µg and each pool of libraries was hybridized to the KAPA HyperExome probes per the KAPA HyperCap Workflow v3.0 user guide but replacing the universal enhancing oligos with MGI blocking oligos. The quality of the pre- and post-enrichment libraries was assessed with an Agilent TapeStation 4200 instrument and the Genomic DNA assay kit (Agilent Technologies). DNA concentration was measured using a Qubit Fluorometer instrument and the Qubit dsDNA Assay Kit (ThermoFisher). Enriched libraries were used for the preparation of DNA Nanoballs. The sequencing was performed on a Complete Genomics G400RS instrument (2×150 bp paired-end sequencing) using standard MPS technology. The average coverage for 4 samples ranged from 154X to 254X, with ≥98% of the regions with a coverage of ≥30x. The fastq read files were aligned to the hg38 genome assembly using bwa (v0.7.171). The duplication detection and removal were performed using Picard (v2.18.11). Copy number profiles for the de-duplicated alignment files for PDX samples were analyzed using Bionano Genomics Inc. Nexus Copy Number 10 software suite build version 9708.

### Analysis of LTL331 Single-cell RNA sequencing (scRNA-seq) data

#### Preprocessing, Quality Control (QC) and Clustering

Using Cell Ranger (v6.0.0), raw sequencing reads were aligned to the reference database combining Ensembl Release 90 of GRCh38 human genome and gene annotation as well as GRCm38 mouse genome and gene annotation. Cells containing > 80 % reads specifically mapping to the human genome were further analyzed, and human-specific reads were used to generate the filtered count matrix and aggregate all 8 samples into a single count matrix using Cell Ranger aggr without normalization. The aggregated count matrix was processed and analyzed further by Seurat (v4.1.1) (24), as follows. First, genes observed in less than 10 cells were discarded. Only cells with (1) mitochondrial transcript percentage below 25%, (2) log-transformed UMI counts and observed genes within 3 median absolute deviation (MAD) from the median, and (3) with at least 1,000 UMIs were kept. Using Seurat v4.1.1 with default parameters, log-normalization, feature selection, and scaling were performed. Principal component analysis was run, and the top 50 PCs were kept based on the knee point selection. Cells were clustered using FindNeighbors and FindCluster functions in Seurat revealing 18 clusters from 51,726 cells. Clusters were visualized by UMAP.

#### Identification of cell states and gene and cell cycle scoring

UCell R package (v2.6.2) (25) was used for scoring AD, NE, EMT and proliferation markers (11) (**Supplementary Table 2**) to annotate the clusters and identify proliferative clusters. The Wilcoxon rank-sum test was used to determine the statistical significance in gene scoring. Each test was repeated three times independently. Cells were assigned to G1, S, and G2/M phases in the cell cycle using the CellCycleScoring function in Seurat.

#### Differentially expressed genes (DEGs) analysis

DEGs of a cluster were determined in comparison to all remaining clusters using the FindAllMarkers function (one-tailed Wilcoxon rank-sum test) in Seurat (**Supplementary Table 3**). P-values were adjusted for multiple testing using the Bonferroni correction.

#### Gene set enrichment analysis (GSEA) and over-representation analysis (ORA)

GSEA was performed to identify significantly enriched pathways in different clusters of single-cell RNA-sequencing data using the hallmark gene sets from the Molecular Signatures Database (MSigDB). The analysis was conducted using several msigdbr (v7.5.1) and fgsea (v1.30.0) R packages. The hallmark gene sets were obtained and prepared by grouping gene symbols by gene set name and converting them into a list structure. GSEA was then performed using fgsea function on gene expression data split by clusters, where a ranked list of genes based on average log2 fold change was analyzed for enrichment. The results were integrated and categorized into three groups as AD, Transition, and NE. A heatmap was generated to visualize the normalized enrichment scores (NES) across clusters. On the other hand, ORA was used to discover enriched pathways using gene sets without ranks such as GEP7. The same packages were used, but enricher function was used to perform ORA instead of fgsea.

#### InferCNV analysis of LTL331 model

InferCNV 2 analysis was performed in R (v4.2.1) using InferCNV library (v1.15.0) (Tickle, Tirosh et al. 2019). All analyzed samples from LTL331/R dataset were subsampled to 2000 cells each unless they had a smaller number of cells to begin with to ensure reasonable processing times and representation in the resulting graph. Luminal epithelial cells from d17 normal adult prostate 3 (26) were used as normal reference. InferCNV analysis was performed for cells grouped by the PDX sample of origin (“sample-level analysis”) and by the clusters identified in our Seurat analysis pipeline. Ultimately, four analyses were performed for cells grouped by clusters and by samples, with cluster_by_groups set to “TRUE” and “FALSE” for each cell grouping type in order to get a better understanding of the distribution of CNV profiles across cell groups. All other settings were common for all analyses: HMM was set to “TRUE” with “i6” HMM type; hclust_method was set to “ward.D2”, noise_filter to 0.01, leiden_resolution to 0.001 and cutoff to the 0.1 (a setting recommended for 10X data).

#### Pseudotime analysis and RNA Velocity in LTL331/R trajectories

We used PAGA to determine the trajectory of 18 clusters and inferred RNA velocity results with scVelo. As a prerequisite, the ‘.loom’ format files that contained matrices for spliced/unspliced counts were generated from FASTQ files of the LTL331 model using 10x Genomics CellRanger pipeline (v7.1.0) and merged into a single file. After that, RNA velocity analysis was carried out using the scVelo python package (v0.2.5) (27). The top 2000 genes with a minimum threshold of 20 expressed counts for spliced and unspliced mRNAs were selected, and pl.velocity_embedding_stream function with default parameters was used initially to annotate clusters with velocity estimations. Velocity pseudotime was determined using tl.velocity_pseudotime function.

#### Random walk analysis and identification of terminal states

We utilized CellRank2 (28) and pyGPCCA (29) python packages to identify the terminal states in the LTL331/R dataset. First, a transition matrix was generated using velocity pseudotime data, obtained as described above. Next, using ‘PseudotimeKernel’ (time key = ‘velocity pseudotime’), we simulated a ‘random walk’ by ‘plot_random_walks’ function to get initial insight into cellular dynamics in the LTL331/R. Generalized Perron Cluster Analysis (GPCCA) estimator (Reuter, Fackeldey et al. 2019) was fitted using ‘fit’ function (n_states = [1, 18]), and the predict_terminal_states function led us to inference of the terminal states, representing stable macrostates. Subsequently, we calculated the fate probabilities of each cluster using the ‘compute_fate_probabilites’ function, which demonstrates the likelihood of transition to terminal states. Since cluster 7 was identified as the sole terminal state among the NE clusters, putative driver genes playing a role in differentiating to cluster 7 were determined by ‘compute_lineage_drivers’ function using a correlation between fate probabilities and gene expression. Lastly, we visualized the expression of putative driver genes along this trajectory based on the velocity pseudotime towards the terminal state cluster 7 using ‘pl.heatmap’ function. The transcription factors used in this analysis were annotated using the transcription factor list from cisTarget database (https://resources.aertslab.org/cistarget/databases/ fetched on August 2022)

#### Single-cell regulatory network inference and clustering (SCENIC)

We utilized pySCENIC (v0.12.0) (30) package to analyze active regulons. Since pySCENIC is nondeterministic, we first subsampled 7,878 cells from the whole dataset and included up to 500 cells per cluster to reduce the computation time. We ran pySCENIC for 20 iterations and only kept transcription factors reported at least 16 times. For each transcription factor, only those targets reported in at least 80% of the runs identifying the transcription factor would be included in the final regulon. This revealed a total of 200 active regulons (**Supplementary Table 4**). Unsupervised clustering of each cell based on its AUC for the 63 regulons ranked as top 5 in at least one cluster was visualized to determine whether AD, NE and EMT clusters can be differentiated based on active regulons. The RSS scores of active regulons can be found in the **Supplementary Table 4**.

#### Public scRNA-seq PCa dataset processing and quality control

Two publicly available PCa scRNA-seq datasets (Referred as ‘Gao dataset’ and ‘Li dataset’) were used (11,12) ()(). All downstream analysis was carried out using the Seurat R package (v4.3.0) (24). Quality control and clustering were carried out as described for the LTL331/R dataset. Patient data from relevant datasets were merged. Batch effects were detected in both datasets using DimPlot function and were corrected using the Harmony integration (v1.1.0) (31). RunHarmony function generated new Seurat objects with Harmony-corrected coordinates to be used in the downstream analysis., Cells were clustered using using FindClusters function with default resolution parameters. clustree R package (v0.5.1) (32) and DimPlot function were used to visualize the clusters. The resolution parameter was selected based on the number of clusters and expression of AD and NE markers. Given the epithelial origin of PCa, we extracted epithelial cells from each dataset. In Gao dataset, epithelial cells were extracted based on the cell type annotation of each cell barcode provided in (11). In Li dataset (12), epithelial cells were annotated using cell marker gene expression. We further confirmed the cell annotation by visualizing the cell marker gene expression. Next, we assigned clusters as AD, NE and “X” based on the expression of lineage markers. The list of cell and lineage markers can be found in **Supplementary Table 2.**

#### Scoring of gene sets derived from LTL331/R dataset

The top 20 differentially expressed genes based on our filtering criteria (log2FC > 0.5, FDR < 0.05) were extracted from each cluster in LTL331/R dataset and were used to score individual clusters from both patient datasets using UCell R package (v2.6.2) (25). The statistical significance of the gene scoring was determined by Wilcoxon-rank sum test between the cluster of interest and other clusters combined.

#### Deriving gene expression programs (GEP) using Consensus NMF

To infer gene expression programs, defined as groups of genes potentially co-expressed or co-regulated under certain conditions in LTL331/R dataset, we applied the cNMF algorithm (v1.4.1) (33). This algorithm takes a count matrix (N cells X G genes) of scRNA-seq data as an input and produces two non-negative low-dimensional matrices, the matrix of GEP (k X G) and the matrix that represents the usage of each program for each cell (N X k). Therefore, we extracted the RNA count matrix from LTL331/R and converted it into ‘float64’ data type and saved it in anndata format. The ‘prepare’ function was initially used to optimize previously generated anndata for running cNMF with default parameters (20 iterations and 2000 highly variable genes), the k parameter values ranging from 5 to 20. After that, the ‘factorize’ function was used to run cNMF, followed by the ‘combine’ and the ‘k_selection_plot’ functions to select the optimal value k based on the stability and error rate. We selected k that showed the most stable solution to run the ‘consensus’ function to compute the consensus solution with density threshold 0.03. The top 100 genes were selected to represent each GEP. Using ComplexHeatmap R package (v2.18.0) (34) we visualized GEPs of LTL331/R in public PCa bulk-RNA sequence datasets.

### Public PCa bulk-RNA seq analysis

We used publicly available bulk RNA-seq patient data (35). Batch correction was carried out as described in the same reference, by using PolyA+ samples as a reference batch for the ‘combat’ algorithm from the sva R package (v3.50.0) (36). Using ComplexHeatmap R package (v2.18.0) (34), batch correction was validated by visualization of AD and NE genes used in the annotation of LTL331/R and compared to clinical annotation.

## Results

### Transcriptional dynamics of cancer cells during NEtD

The LTL331 model provides a unique framework for studying NEtD (14,15) as this model begins as AR-driven PRAD and undergoes NEtD following host castration over approximately 20-22 weeks (15). To gain a better understanding of the NEtD, we analyzed tumor samples collected at defined time points, including pre-castration (preCx) and at 2, 4, 8, 12, and 16 weeks post-castration (postCx), as well as from two relapsed tumors at 20 and 22 weeks. Tumor response to host castration was evident by substantial decrease in PSA levels by 2 weeks postCx and reached to baseline by week 4. In parallel, tumor volume decreased by approximately 75% by week 4 and continued to decrease until relapse at week 20 (**Fig. 1A**). AR protein expression was detectable during the postCx period; however, PSA expression was absent by week 4 postCx and more importantly, NEPC markers were not detected until 16 weeks postCx (**Fig. 1B**), indicating a temporal lag between AR loss and emergence of NEPC features (**Fig. 1B**), in line with previous observations. These samples were subjected to whole exome sequencing (WES) and copy number variation (CNV) analyses to assess the genomic stability and compare to previous generations of LTL331 and donor patients. CNV profiles were consistent with previously determined and published CNV profiles for the parental tumor and earlier generations of the LTL331 model (**Supplementary Fig.1-2**). Furthermore, InferCNV profiles of the LTL331 model suggests that the majority of CNV features remained conserved across all timepoints and phenotypic states (**Supplementary Fig.5-6**), reinforcing the conclusion that NEtD is driven by transcriptional reprogramming (14,15).

**Figure 1.**
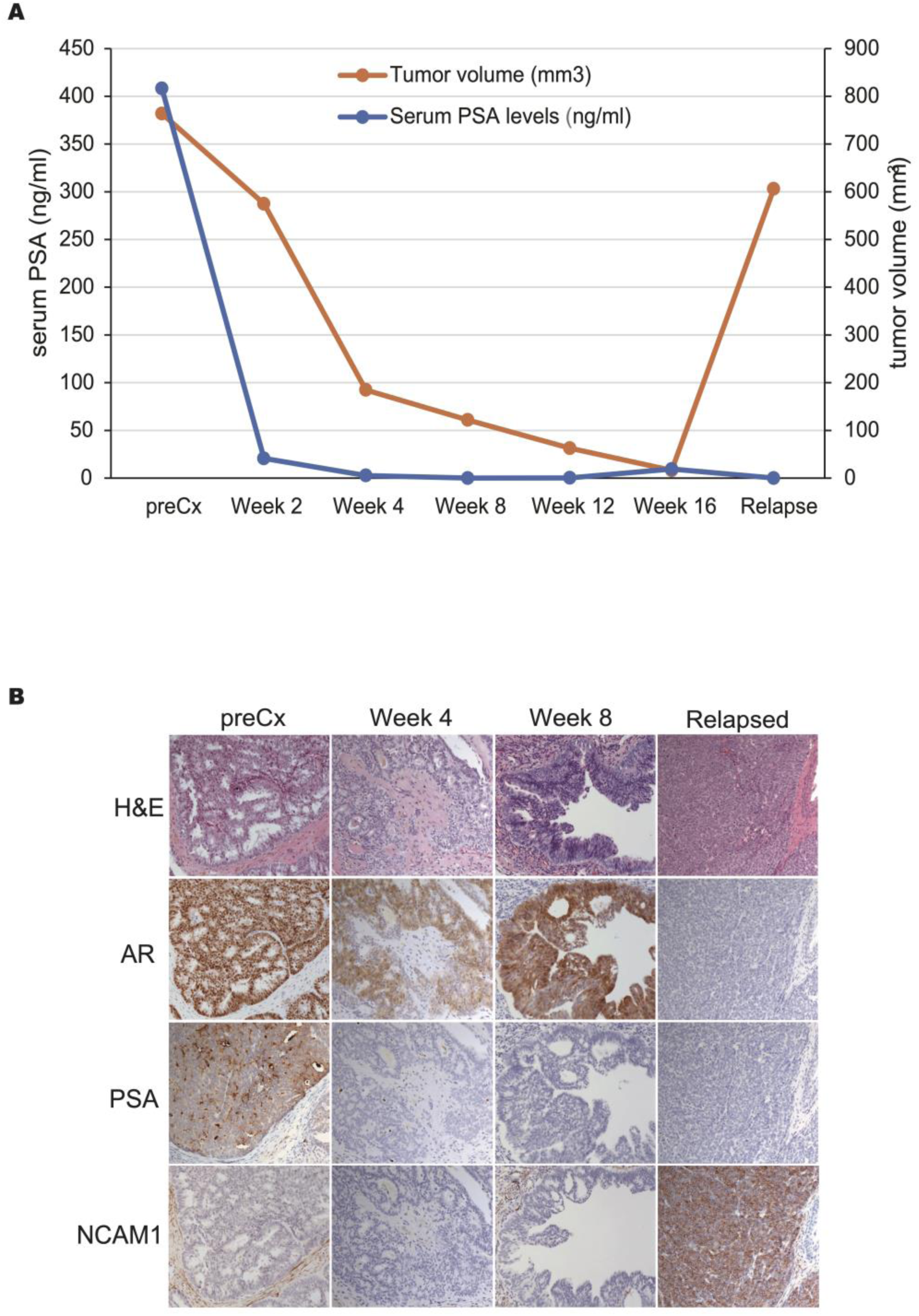
Time course of the development of LTL331 model. **(A**) PSA level and tumor volume change across selected time points in the development of LTL331 model. **(B)** H&E staining of *representative* LTL331 tumors for selected time points and IHC analysis of expression of key proteins characteristic for PCa and NEPC.

From the remaining tumor, we obtained 51,726 high-quality single-cell transcriptomes (**Supplementary Table 1**). Unsupervised clustering revealed distinct expression patterns across 18 clusters (**Fig. 2A**), including 10 clusters characterized by varying degrees of AR, one intermediate cluster with low AR and NE markers (AR-low/NE-low), and seven clusters predominantly neuroendocrine with high NE marker expression and negligible AR expression (AR-/NE+ NEPC) (**Fig. 2B and C**).Two preCx PRAD clusters 11 and 15 demonstrated the highest AR, KLK2, KLK3, and TMPRSS2 expression, consistent with high AR activity (**Fig. 2C and D**). We note that these clusters do contain a small number of cells (∼ 2%) with undetectable AR axis activity. In contrast, the other eight postCX PRAD clusters exhibited lower AR activity supported by low KLK3 expression. All seven NEPC clusters were characterized by positive NE scores, no AR expression and very low to no AR activity, consistent with their NE phenotype. Notably, cluster 17 represents an intermediate state with diminished PRAD characteristics (lower AR expression and activity) and NE lineage markers (e.g., NCAM1) (**Fig. 2C and D and Supplementary Table 2**). Further analysis of epithelial markers showed consistent expression of luminal cytokeratins (KRT8, KRT18) across all PRAD and NEPC clusters, while basal cytokeratins (KRT5, KRT14) were absent (**Fig. 2E**). These findings reinforce that NEPC development involves lineage reprogramming from a luminal origin and further shows that LTL331 model captures tumor heterogeneity and cellular lineage states during NEtD.

**Figure 2.**
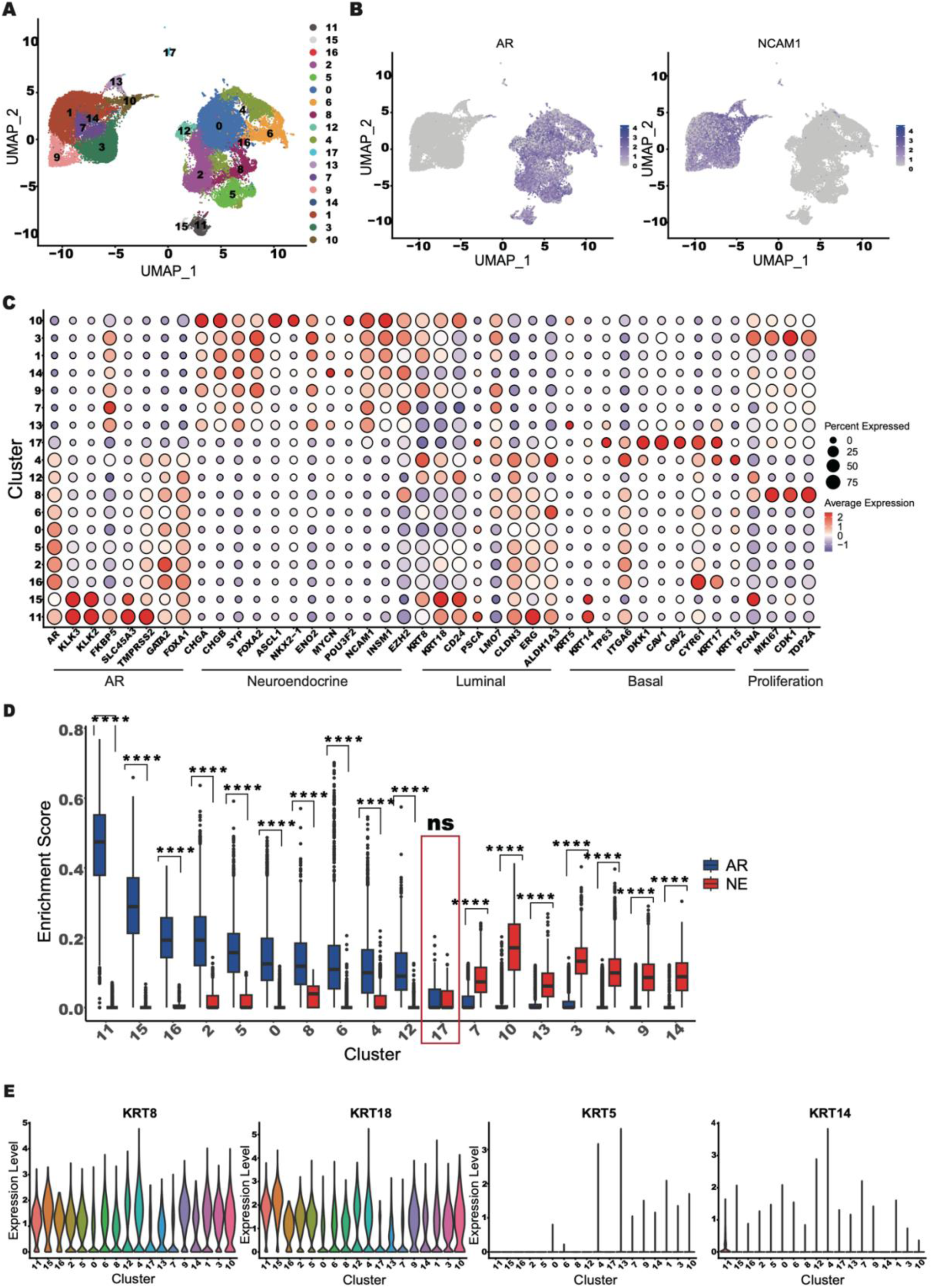
Transcriptional heterogeneity of cancer cells in NEtD. **(A)** UMAP visualization of 51726 transcriptomes colored by clusters. **(B**) UMAP visualization of AR and NCAM1 gene expression. **(C)** Dot plot displaying the expression of canonical lineage markers and signature genes, grouped by type and displayed at the bottom. The size of the dot represents the percentage of cells in the cluster expressing the gene and color shows the level of expression. **(D**) Violin plot showing enrichment scores for PRAD (blue) and NE (red) marker genes (**** represents p-value smaller than 0.0001, ns denotes not significant). **(E)** Violin plots showing luminal and basal cytokeratin gene expression.

### Longitudinal analysis revealed dynamic cell state changes during NEtD

To investigate how cellular states evolve during NEtD, we next characterized changes in cellular composition as tumors transitioned from PRAD to NEPC following castration. Our longitudinal analysis of scRNA-seq data demonstrated that castration in this unique model induced significant and dynamic alteration in cellular states, marking a progression from PRAD to NEPC (**Fig. 3A and B**). Initial analysis of the preCX tumor revealed substantial intra-tumoral heterogeneity (**Fig. 3A and B**), with eight out of ten PRAD clusters present (**Fig. 3A and B**). Clusters 11 and 15 were predominant, comprising 63% and 15% of the tumor, respectively (**Fig. 3A and B**), with the remaining six clusters (0, 2, 4, 6, 8, and 12) contributed minor fractions (ranging from 0.4% to 13%). Though some of these persisted in the relapsed tumor, altogether they constituted only 0.2% of the tumor mass (**Fig. 3A and B**), highlighting the enduring heterogeneity as the tumor evolves towards NEPC. Crucially, NEPC-specific clusters were absent in the preCx tumor and predominantly appeared in later postCx phases, indicating a shift in cell lineage from PRAD to NEPC, rather than an expansion of pre-existing NE cells (**Fig. 1B**).

**Figure 3.**
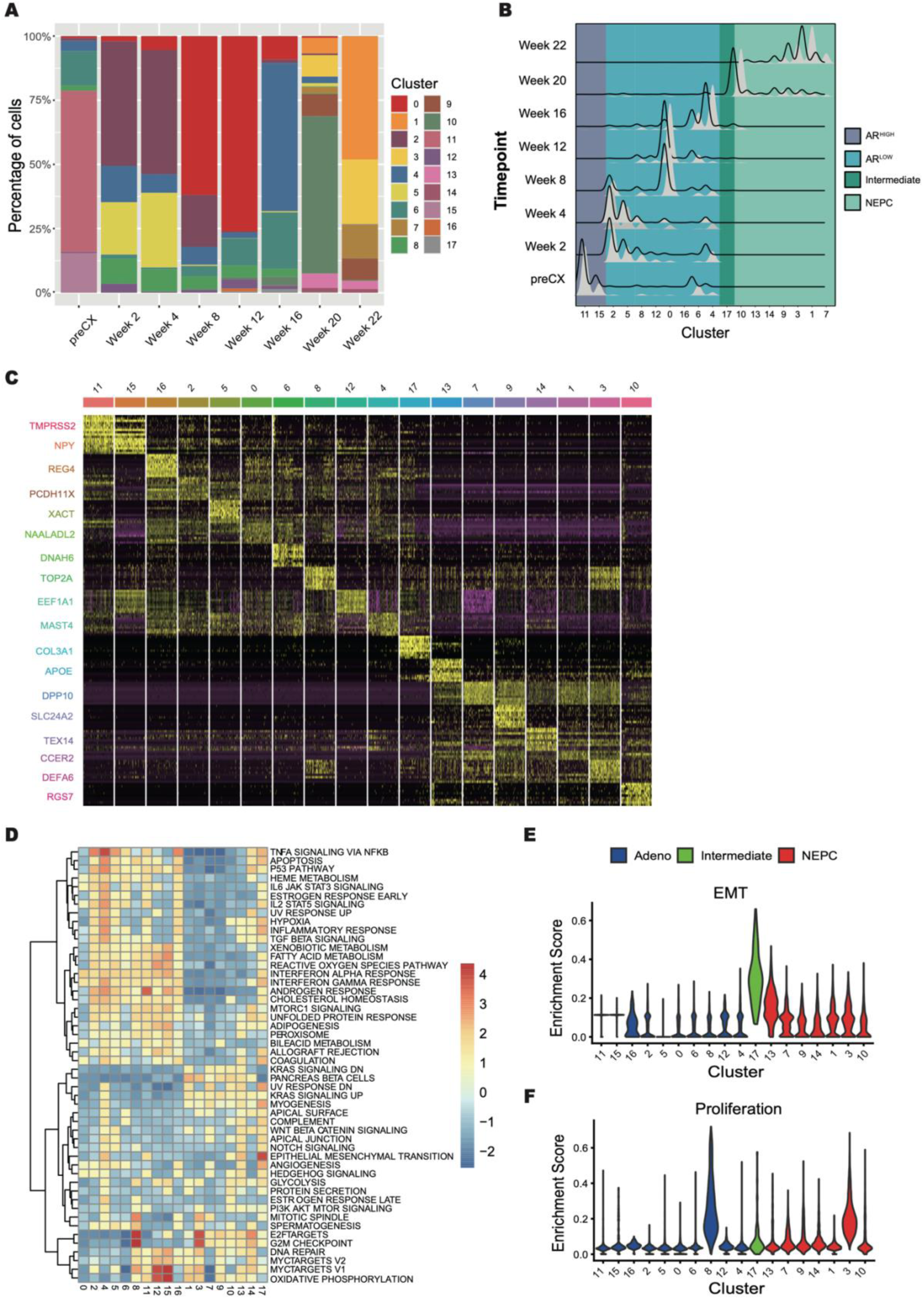
Longitudinal analysis demonstrated a dynamic change of cancer cell state during NEtD. **(A)** Bar graph displaying the relative proportion of clusters in LTL331 dataset across time points. **(B)** Ridge plot displaying the number of cells in each time point scaled to even out representation of cells across time points. **(C)** Heatmap showing the top 15 differentially expressed genes identified by multi-pairwise comparison of all clusters. Select genes are highlighted. **(D)** Heatmap showing gene set enrichment of hallmark pathways. **(E)** Violin plot showing enrichment scores for EMT related genes or Bar graph showing gene set enrichment of EMT, progenitor and stem cell related gene sets in cluster 17. **(F)** Violin plot depicting enrichment scores for proliferation related genes.

We categorized the dynamic shifts in tumor composition into the four distinct response groups. (1) AR^high^ PRAD clusters (11 and 15): Dominated the preCX tumor (80%) but were undetectable two weeks postCx (**Fig. 3A and B**). Despite the loss of these two AR^high^ clusters, the tumor volume only decreased by ∼25% at this time point (**Fig. 1A**), suggesting a rapid adaptation of their transcriptional states that contribute to later-stage clusters. (2) AR^low^ PRAD clusters (0, 2, 4, 5, 6, 8, 12, and 16): Initially low in AR expression/activity, these clusters underwent a significant, albeit transient, expansion postCx (**Fig. 3A and B**). Consistent with recent reports the number of AR and AR activity low cells increased from ∼15% to ∼100% (37). Clusters 8 and 12 exhibited relative stability, with only brief expansion two weeks postCx. (3) Intermediate cluster (17): Emerged at 8-12 weeks postCX. (4) NEPC clusters (1, 3, 7, 9, 10, 13, and 14): Absent in the preCX tumor, these clusters emerged in later phases, mainly 20 weeks postCX, with cluster 10 appearing earlier at 8-12 weeks postCX, indicating a PRAD-NEPC lineage shift over pre-existing NE cell expansion.

While multiple clusters expanded simultaneously, only one became dominant at any given time, constituting 50% or more of the tumor. Sequentially, PRAD clusters 11 (preCx), 2 (2 and 4 wks), 0 (8 and 12 wks), 4 (16 wks), and NEPC clusters 10 (20wks) and 1 (22wks) represented the dominant transcriptional states, marking the transition from AD to NEPC (**Fig. 3A and B**). To define the molecular characteristics of these transcriptional states, we analyzed differentially expressed genes (**Fig. 3C and Supplementary Table 3**) and conducted a gene set enrichment analysis (GSEA) using hallmark pathways and chemical genetic perturbation (C2) gene sets from the Molecular Signatures Database (MSigDB). Several hallmark gene sets, such as P53 pathway, TNFα signaling, interferon alpha, and androgen response, were enriched in both AR^high^ and AR^low^ PRAD clusters, whereas others like pancreas beta cells and E2F targets were prevalent in NEPC clusters (**Fig. 3D**). Specific pathways such as notch signaling in NEPC cluster 13 and TNFα signaling and EMT and TGF-β signaling in PRAD cluster 4 distinguished these clusters from others. The EMT, myogenesis, and angiogenesis pathways were particularly enriched in EMT were also differentially expressed in this cluster (SNAI2, ZEB1/2, TWIST1/2). In addition, this cluster displayed enrichment for stem and progenitor cell-related gene sets (**Fig. 3E**). In parallel, we observed a large subset of genes important for prostate luminal progenitors, prostate cancer stem cells being expressed in clusters 4, 17 and 13 (**Supplementary Fig. 3A)**. The expression of neural crest stem cell genes, including CD44 (38) and THY1 in cluster 17 is intriguing **(Supplementary Fig. 3B)**, as caudal neural crest cells were suggested (39).(39). Additionally, cluster 17 also expresses several known mesenchymal stem cell markers, including NT5E/CD73, ITGA1/CD29, CD44 (40), and STRO-1/CD34 (38). Collectively, this implies that cells in this cluster are going through dedifferentiation and gaining stem cell-like characteristics.

To determine whether proliferative capacity drives cluster expansion, we assessed proliferation scores using canonical markers such as PCNA and KI67 (**Fig. 3F and Supplementary Table 2**). Although clusters 8 (PRAD) and 3 (NEPC) scored higher than others, neither cluster dominated at any stage. Additionally, no discernible differences in cell cycle profiles were observed before or during expansion (**Supplementary Fig. 4**), suggesting that proliferative capacity alone does not account for the observed changes in relative cluster sizes. Instead, dynamic changes in cluster sizes imply modifications in their transcriptional states, reflecting changing levels of cell adaptation to postCx conditions.

### Epithelial-mesenchymal transition highlights the transitional cluster during NEtD in the inferred trajectory

To gain insight into NEtD, we carried out partition-based graph abstraction (PAGA connectivity analysis (41) to capture the proximities of clusters at either end of the phenotypic range (**Fig. 4A**). Using AR^high^ cluster 11 as the root, connectivity analysis revealed that the EMT stem like cluster 17 forms a key transition point in NEtD. This is in line with its intermediate state with low AR and NE signature scores, timing of its emergence, expression of EMT, and stemness genes.

**Figure 4.**
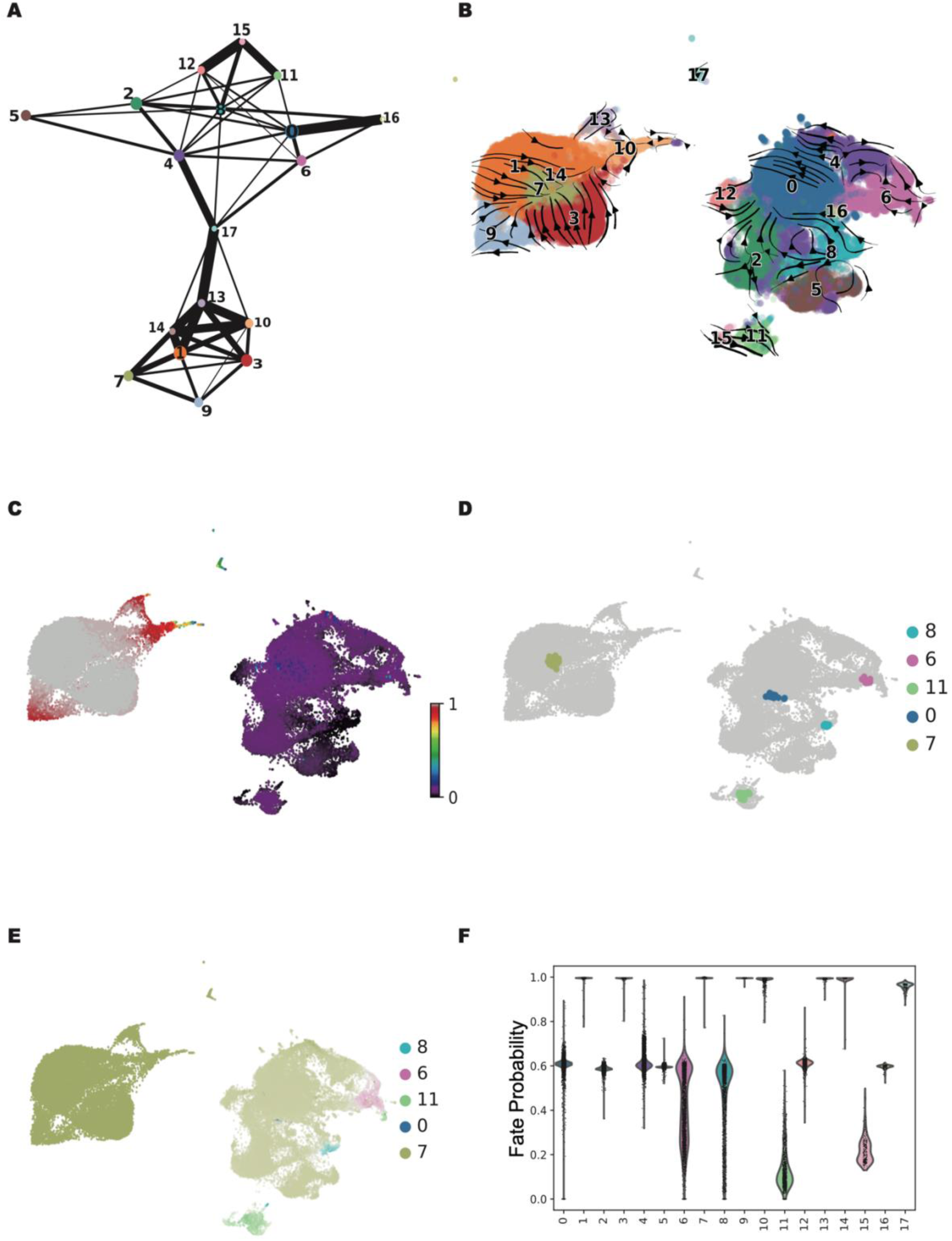
Epithelial mesenchymal transition highlights the transitional cluster during NetD in the inferred trajectory. **(A)** PAGA connectivity map. Each node displays a cluster. The edges between the clusters show the connectivities. The connectivities with weights higher than 0.3 are shown. **(B)** RNA velocity embedding on UMAP plots. Arrows point towards the predicted course of cell state changes. **(C)** Visualization of cells colored by RNA velocity pseudotime. **(D)** Terminal states during NEtD. **(E)** Fate probabilities projected onto UMAP. Cells are colored based on the identity of the terminal states and the intensity of the color reflects the degree of fate probabilities. **(F)** Violin plot showing fate probabilities of cells in individual clusters towards terminal state cluster 7.

PRAD clusters 4 and 6, which exhibited similar patterns of expansion reduction prior to mass emergence of NEPC clusters, showed strong connections to cluster 17 (**Fig. 4A**), suggesting that both clusters might transition into EMT stem like state. The path of transition bifurcated after cluster 17 to clusters 13 (ASCL1 negative) and 10 (ASCL1 positive). While cluster 13 is classified as a NEPC cluster, its relatively low NE gene signature score compared to other NEPC clusters suggests that it represents an early NEPC state (**Fig. 2D**). The strong connectivity between clusters 10 and 13 implies a potential transitional relationship between these two states (**Fig. 4A**). Interestingly, both clusters converge onto cluster 14, indicated by their strong connections (**Fig. 4A**), suggesting that this cluster may represent a subsequent stage of NEtD. Notably, both proliferative clusters 3 (NEPC) and 8 (PRAD) exhibit the highest number of connections (**Fig. 4A**), raising the possibility that these populations may contribute to replenishment of other cellular states.

To infer directionality in these state changes we next estimated RNA velocities using scVelo as it allows dynamic modeling of transcription and splicing kinetics. RNA velocities were largely in line with PAGA results. For example, cells in AR^low^ cluster 4 have three trajectories, pointing to cells in cluster 6 (AR^low^ PRAD),17 (intermediate EMT stem like), or 10 (NEPC), while cells in AR^low^ cluster 0 are mainly pointing toward cluster 10 (**Fig. 4B**). RNA velocities provided directionality to the connectivities between several AR^low^ clusters. For instance, cells in several clusters, including 2, 12, 5 and proliferative cluster 8 point towards clusters 0 or 4, suggesting that the increase in size of these two clusters at the intermediate time points might result from the state transitions (**Fig. 4B**). When we examined the trajectories of the NEPC clusters, we observed several clusters’ trajectories point towards cluster 7. Pseudotime analysis identified the EMT stem like cluster 17 as focal point of NEtD (**Fig. 4C**). It also positioned PRAD clusters 4 and 6 as pre or early EMT states, and NEPC clusters 10 and 13 as late EMT/early NEPC (**Fig. 3D and 4C**). These findings align with EMT hallmark enrichment and EMT regulator expression in clusters 4 and 13, although at lower levels than in cluster 17 (**Fig. 3D and Supplementary Fig. 3A**).

### Transcriptional programs in NEtD

To understand gene regulatory networks (GRN) underlying NEtD, we performed single-cell regulatory network inference and clustering (SCENIC) analysis (30). High-confidence active regulons, consisting of transcription factors and their target genes, were defined based on the consensus of twenty independent runs, retaining regulons in at least 80% of the iterations. The hierarchal clustering of active regulons distinguished EMT stem like, PRAD, and NEPC clusters, including FOXA1 and NKX3.1 regulons for PRAD, FOXA2, ASCL1 regulons for NE and MSX1, RUNX3 regulons for EMT. Interestingly, both highly proliferative clusters 3 (NEPC) and 8 (NEPC) were marked by FOXM1, E2F1/2/8 and MYBL1/2 regulons (**Fig. 5A and Supplementary Table 4**).

**Figure 5.**
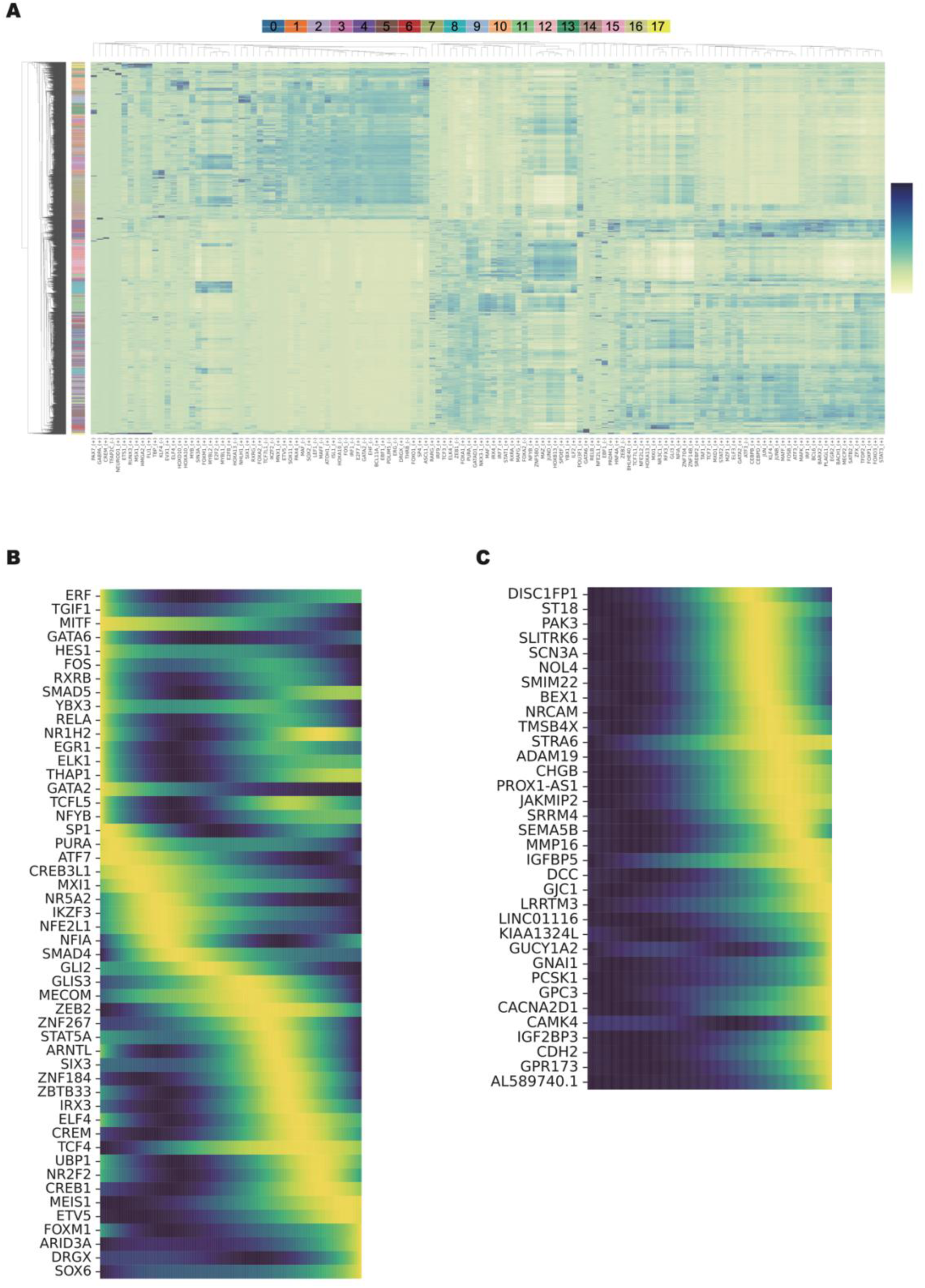
Transcriptional programs in NEtD. **(A)** Active regulons differentiating PRAD, NE and EMT clusters. **(B)** Smoothened gene expression trends of transcription factors in the putative driver genes predicted by CellRank2 along the pseudotime towards cluster 7. **(C)** Smoothened gene expression trends of the top 50 putative driver genes predicted by CellRank2 along the pseudotime towards cluster 7.

To identify TFs specifically involved in NEtD, we further focused on the top 20 regulons active within clusters central to the transition, including clusters 4, 6, 17, 10 and 13, which represent early to late EMT states based on pseudotime analysis (**Supplementary Fig. 7**). IRF1, SOX9, and KLF5 were the top regulons of cluster 4, which is representative pre/early EMT cluster. Notably, KLF5 has been shown to antagonize AR-driven transcriptional programs (42) and is required for TGF-β -induced EMT in PCa cell lines (43). SOX9 overexpression also enhances NE-like features in PCa cell lines (44), although its role in NEtD has not been fully explored. In cluster 6, RFX2 and RFX3, both regulators of multiciliated cell differentiation, suggest an enrichment of ciliogenesis in this population. Other top regulons in cluster 6, including FOXA1, ZEB1, and ZEB2, are linked to EMT and prostate cancer plasticity, while FOXP1 is associated with neural development. Together, these features align with characteristics of pre- or early EMT cell populations.

Cluster 17 exhibited the higher number of uniquely active regulons, including MSX1, RUNX3, NEUROD1, GABPA, HMGA1/2, CREM, TCF7L1, DLX2, and RELB. Of particular interest, TCF7L1, a factor involved in stem cell pluripotency (45) and brain development (46), has recently been implicated in promoting NEPC features under androgen deprivation(47). Analysis of active regulons revealed that this cluster could be further subdivided into early and late regulatory states, sharing transcriptional programs with PRAD and NEPC clusters respectively (**Fig. 5A**). Among these regulons MSX1 was shared between PRAD cluster 4 and EMT stem like 17, showing robust expression and high activity in cluster 17, but lower expression in cluster 4 (**Supplementary Fig. 3B**). This suggests MSX1 may serve as an early driver of EMT, consistent with its role in neural crest specification (48,49).

Cluster 10 was characterized by the activation of several key regulons, including HOXD10, ASCL1, HOXA10, ELF4, SP4, PGR, and KLF12 (**Fig. 5A and Supplementary Fig. 7**). Among these, ASCL1 is a well-established NE lineage TF in the lung and a known driver of SCLC (50). ASCL1 is highly expressed in NEPC patients (51,52) and PDX models (51,53) and was shown to activate neuronal and stem-like programs in PRAD cell lines (54) and to play a critical role in NEtD (53). In contrast, SP4, HOXA10, HOXD10, KLF12, and EBF4/MEF have not been previously implicated in NEPC.

Cluster 13 was instead marked by distinct active regulons namely PAX7, RXRG and FOXG1 **(Fig. 5A and Supplementary Fig. 7)**. RXRG (55–57) and FOXG1 (58,59) are important regulators of nervous system development. Notably, PAX7 is required for neural crest specification during embryogenesis (60) and is expressed in progenitor populations of neural crest descendants (61), suggesting a potential link between neural crest programs and early NEPC development.

### Cell Fate and Potential Lineage Drivers for NEtD

To further elucidate cellular fate transitions and terminal states during NEtD, we applied CellRank2 (28), which infers terminal states, estimates fate probabilities for each cell, and identifies putative driver genes associated with these terminal states (28,29). Using CellRank2, we identified four terminal states of cells in both AR^high^ and AR^low^ PRAD cluster, specifically in clusters 11, 0, 6, 8, and one terminal state in NEPC cluster 7 (**Fig. 4D**). The presence of distinct terminal states within PRAD clusters is particularly intriguing, as it highlights substantial heterogeneity: while certain cells retain capacity to transition between states, others have already reached a terminal differentiation stage. This indicates that LTL331 enables the captures of tumor heterogeneity across and within clusters, offering enhanced resolution of lineage trajectories that bulk RNA-seq data often misses. Notably, most cells exhibited fate bias towards NEPC cluster 7, suggesting that regardless of the NEtD path taken, cluster 7 represents the ultimate terminal state (**Fig. 4E and F**).

We then extracted lineage driver genes involved in cellular transitions toward this terminal NEPC state. Putative drivers were identified based on systematic upregulation (or downregulation) in cells with higher (or lower) probability of differentiating into cluster 7. From these candidates (**Supplementary Table 6**), we prioritized transcription factors (TFs) for further examination (**Fig. 5B**). Among those, TCF4 is a notable TF previously implicated in NEtD in both PCa (62) and lung NE cancer (63). Another TF, ST18 (MYT3) is highly expressed in NEPC (64,65), small cell lung cancer (SCLC) (65), and small intestine neuroendocrine tumors (66). Its paralog, MYT1L, along with ASCL1 and POU3F2, has been shown to play crucial roles in reprogramming fibroblasts and hepatocytes into functional neurons (67,68). SP1 emerged as another critical factor. It modulates multiple signaling pathways essential for EMT, including TGF-β (69), the Wnt/β-catenin (70), and possibly the Notch pathway (71). SP1 also regulates key EMT TFs such as Snai1 and Snai2, upregulated in cluster 17 (72). Importantly, SP1 has been shown to be druggable in preclinical models of castration resistant PCa including NEPC models. In particular, an FDA-approved small molecule plicamycin demonstrated significant antitumor activity both *in vitro* and *in vivo* in these models (73).

In addition to TFs, we also examined non-TF putative driver genes as they may contribute to NEtD through alternative regulatory or functional mechanisms (**Fig. 5C**). For instance, SRRM4 plays a role in NEtD by regulating the alternative splicing of REST, a repressor of neuronal gene expression, compromises its repressive capacity (22,74,75). Several other candidate genes had not been linked to NEtD previously. For instance, SCN3A encodes a voltage-gated sodium channel and is expressed early in nervous system development, including in cortical progenitors and neurons (76,77). The long noncoding RNA PROX1-AS1 1 has been implicated in various cancers, although its role in NEtD remains unexplored (78–80). Recently, SP1, which increases in expression early during NEtD (**Fig. 5B**), has been shown to be active in colorectal cancer as an upstream activator of PROX-AS1 (81). Together, these findings reveal a transcriptional roadmap of NEtD, delineating a progression from PRAD clusters through an EMT stem like intermediate toward NEPC states. Along this trajectory, we identified putative regulators at each stage, including KLF5, TCF4, TCF7L1 as potential early EMT stemness drivers, ASCL1, PAX7 and RXRG as candidate fate determinants. These results provide opportunities to further dissect the regulatory networks and lineage plasticity underlying NEtD.

### Temporal Transcriptional programs in LTL331 confirmed in clinical data

To evaluate the clinical relevance of our findings, we assessed whether distinct temporal populations observed in LTL331 model could be identified in publicly available scRNA-seq datasets from PCa patients. We first analyzed a dataset (Gao dataset) comprising samples from one prostatic intraepithelial neoplasia, two PRAD, and three NEPC patients(11). The dataset was processed using our analysis pipeline (**Supplementary Fig. 8A**), focusing specifically on the luminal epithelial cells (**Supplementary Fig. 8B and C**). Based on the expression of AR pathway and NE markers, we annotated the clusters as PRAD (Gao_0, Gao_17, and Gao_18), NEPC (Gao_8, Gao_11, and Gao_13) and intermediate (Gao_4, Gao_14, Gao_16, and Gao_20) populations (**Supplementary Fig. 8B and C**). We systematically scored them using LTL331 cluster-specific genes (**Fig. 6 and Supplementary Fig. 9**). The similarity between the PRAD clusters were primarily driven by the level of AR expression and activity. For instance, patient-derived clusters Gao_0, Gao_17 and Gao_18 showed the highest similarity scores when assessed with gene signatures from LTL331 clusters 11, 15 or 2 (**Fig. 6A-E**), all of which are characterized by high AR activity and AR expression (**Fig. 2D**).

**Figure 6.**
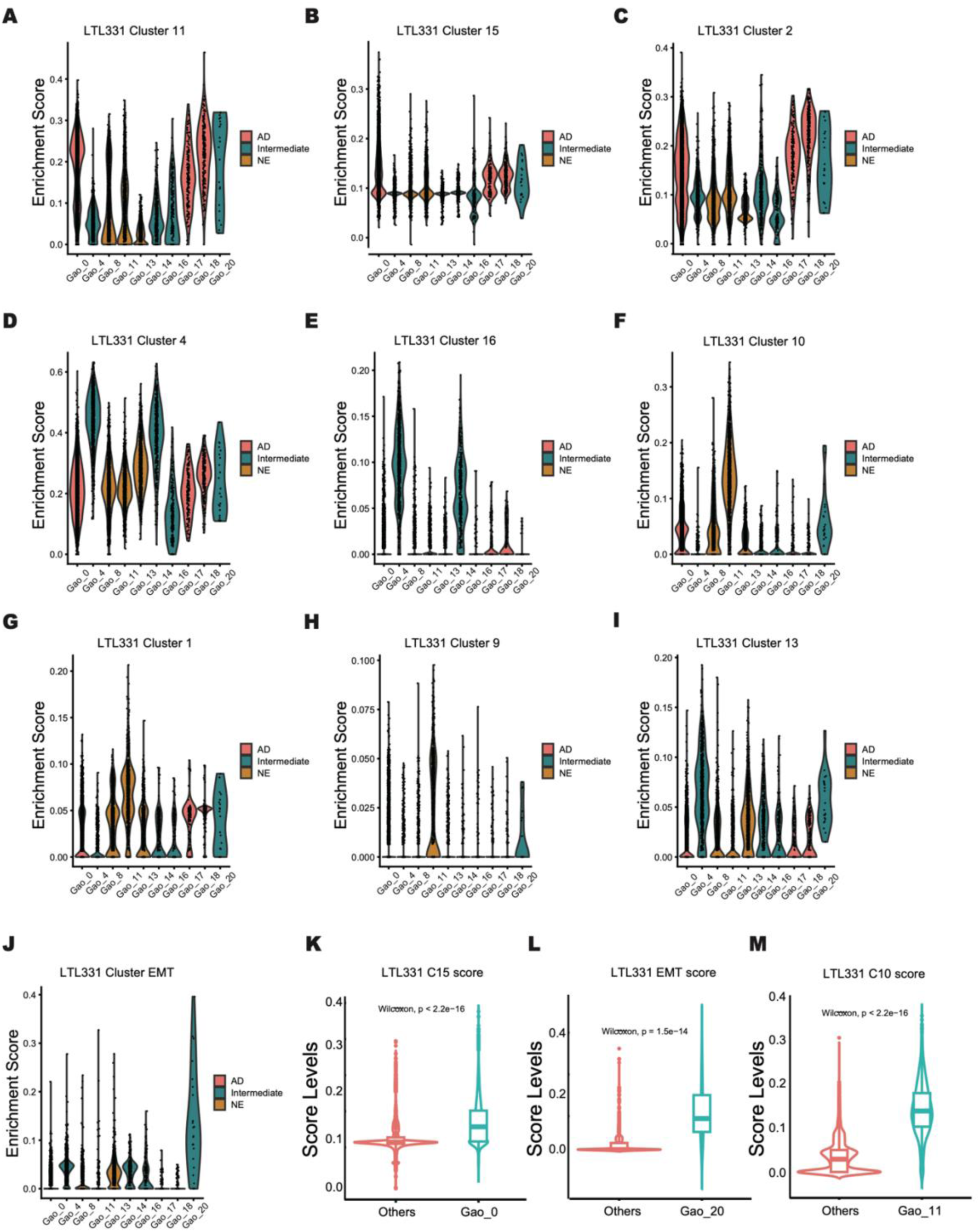
Transcriptional programs in LTL331 dataset confirmed in clinical data. **(A-E)** Violin plots showing enrichment scores of select clusters for PRAD cluster-specific genes**. (F-J)** Violin plots showing enrichment scores of select clusters for NE and EMT cluster specific genes. **(K-M)** Violin plots depicting enrichment scores of patient clusters 0, 20 and 11 using PRAD, NEPC or EMT cluster-specific genes and others (Wilcoxon test, the boxplot represents the interquartile range (IQR) divided by the median).

Gao_11 (NEPC) from the patient dataset exhibited strong similarity to cluster 10 in LTL331, likely driven by ASCL1 expression, a key TF associated with NEtD (51–54). ASCL1 was robustly expressed in both clusters, supporting the alignment of their transcriptional programs (**Fig. 6F**). Additionally, Gao_11 also showed partial similarity with LTL331 clusters 1 and 9 (**Fig. 6G and H**), suggesting these clusters may still have residual ASCL1 transcriptional programs. Gao_4 and Gao_20 (Intermediate) from this dataset scored highly when evaluated with gene signatures from LTL331 cluster 13 (**Fig. 6I**), representing a late EMT/early NEPC state. Importantly, the gene signature of LTL331 cluster 17 (**Fig. 6J**), representing a population central to NEtD, was also detected in Gao_20, suggesting that this transitional state is not model-specific but also occurs in human tumors. To further support the clinical relevance of these findings, we determined the fraction of cells contributed by individual patients to each of these clusters (**Supplementary Fig. 8D**). The presence of LTL331-like populations across multiple patients strengthens the conclusion that these cellular states are bona fide components of the NEtD trajectory in clinical NEPC.

To further validate the clinical relevance of our findings, we analyzed a second set (Li dataset) of scRNA-seq data (12), comprising cells from two hormone-sensitive and two castration-resistant PCa specimens and identified five PRAD (Li_0, Li_5, Li_8, Li_10, and Li_16) and three NEPC (Li_1, Li_4, and Li_9) clusters based on AR pathway and NE markers gene expression (**Supplementary Fig. 10A-C**). While most clusters contained cells derived from multiple patients, Li_0 was unique to a single patient (**Supplementary Fig. 10D**). Importantly, Consistent with our initial analysis, AR activity, AR or ASCL1 expression remained the key determinants in identifying analogous to those found in the LTL331 model (**Supplementary Fig. 11 and 12**).

We next investigated whether transcriptional programs identified in the LTL331 model via scRNA-seq could also be detected in bulk RNA-seq datasets from PCa patients. Using consensus non-negative matrix factorization on LTL331 scRNA-seq data, we consolidated 18 clusters into seven gene expression programs (GEPs) (**Supplementary Table 7**), comprising two NEPC-specific programs, four PRAD-specific programs, and one proliferation-associated program shared by highly proliferative cells.

To evaluate the clinical relevance of these GEPs, we analyzed a large collection of bulk RNA-seq dataset comprising 316 CRPC and 19 NEPC samples (35). Patients were first stratified into four groups (AR+/NE-, AR-low/NE-low, AR-/NE+, AR-/NE-) based on AR pathway activity and NEPC marker gene expression **(Supplementary Fig. 13A)**. Hierarchical clustering using the top thirty genes from each GEP revealed that both PRAD- and NEPC-associated GEPs were readily detectable in clinical samples **(Supplementary Fig. 13-15)**. This supports that transcriptional states analogous to those in the LTL331 model are present across CRPC and NEPC patient tumors.

## Discussion

LTL331 model exhibits highly reproducible phenotypic trajectory, consistently initiating as PRAD and undergoing transdifferentiation into NEPC upon host castration. This predictable lineage progression offers a robust model to dissect the molecular underpinnings of NEtD in prostate cancer. The genomic stability of LTL331 model, supported by consistent copy number variations across generations and with the original donor patient and the expression of luminal cytokeratins in the relapsed tumor further validates its utility for studying NEtD and supports the idea that NEPC cells originate from transdifferentiating luminal PRAD cells (11,82).

Longitudinal single cell transcriptomic profiling across the NEtD trajectory unveiled a continuum of PRAD to NEPC states. This transition was characterized by the gradual loss of clusters with high AR activity and the concurrent acquisition of neuroendocrine features. Castration-induced transcriptional reprogramming drove dynamic shifts in cellular populations, marked by cyclical expansion-reduction and changes in the states of key clusters such as AR^low^ clusters 4 and 6. These shifts culminated in the appearance of an EMT/stem-like cluster 17, characterized by transcriptional features of de-differentiation and stemness. Pseudotime analyses positioned cluster 17as a transitional node linking AR^low^ PRAD states to early NEPC states. Our single-cell analysis further extends beyond the identification of EMT transition states to establish a more comprehensive model of the NEtD mechanism. We show that pre-EMT cells, enriched for ciliogenesis and cell adhesion pathways, differentiate into EMT-like populations. These EMT-like cells then progress into distinct ASCL1+ and ASCL1− NEPC states, revealing a hierarchical trajectory that underlies therapy resistance. The key findings at each stage of this progression are outlined below.

### EMT and Neural Crest Stem Cell States

The identification of EMT stem-like cluster 17 as a pivotal transitional node between PRAD and NEPC aligns with the established role of EMT in the relapse of NEPC (15,83). This cluster is enriched in canonical EMT master regulators (ZEB1/2, TWIST1/2, SNAI2) and EMT-related active regulons, including HMGA1 (84) and MSX1 (85). Interestingly, MSX1, a key driver in neural crest specification (85), induces a dedifferentiated progenitor-like state when ectopically expressed in melanocytes and myotubes (85), resembling its role in cluster 17. Similarly, GLI2, which is critical for induction and migration of the neural crest in *Xenopus laevis* (86), is also highly expressed in this transitional population. Significantly, these cells exhibit enrichment of progenitor and neural crest stem cell gene signatures (e.g., LEE_NEURAL_CREST_STEM_CELL_UP), suggesting concurrent EMT activation and gaining stem cell-like characteristics during NEtD. While prior studies have hinted at nerual crest-like progams in NEtD (87), our work provides the strongest evidence to date linking these pathways to disease progression. This connection is further supported by the neural crest’s role in normal prostatic neuroendocrine cell development (82), implying conserved mechanisms in pathological NEtD. These findings position EMT/stem-like cluster 17 as a critical therapeutic target to disrupt NEtD. Future studies should clarify how MSX1, GLI2, and associated regulators orchestrate this transition and identify druggable nodes within neural crest-related pathways.

### Ciliogenesis Signature in pre/early EMT cells

Understanding how late-stage PRAD clusters (0, 4, and 6) transition into EMT/stem-like and early NEPC-like states is critical for developing therapies to prevent NEtD. These clusters are detectable in pre-castration tumors and show transient expansion despite overall tumor shrinkage, offering a potential window for early intervention. While multiple tumor clusters express prostate progenitor and stem cell-related genes, clusters 4 and 6 exhibit a broader spectrum of markers, indicating the initiation of de-differentiation within these populations. In line with this, cluster 4 and 6 show enrichment of several pathways known to induce EMT, including JAK/STAT (88), hypoxia (89) and TGF-β (90). Further ORA analysis revealed that cluster 6 is enriched for ciliogenesis and ciliopathy-associated pathways (**Supplementary Fig. 16B**) (91), particularly within a small terminal subpopulation identified by CellRank2. Comparison with cluster 17 showed that cluster 6 harbors a distinct ciliogenesis signature (**Supplementary Fig. 17A**), potentially indicative of motile cilia. Given the context-dependent role of cilia in tumor progression, including dormancy regulation through cell cycle control and metastasis via microenvironmental signaling, this finding suggests a novel functional state in prostate cancer (92,93). Significantly, this ciliogenesis signature was also observed in RNA-seq data from PDX models of cellular dormancy (**Supplementary Fig. 17B**) and in a clinical cohort (**Supplementary Fig. 16A**) (87). Together, these results suggest that cluster 6 subpopulation may represent a quiescent, persistent state (92).

### ASCL1-independent Pathways in continuum of NEPC Phenotypes

Following the EMT/stem-like state, the trajectory toward NEPC bifurcates into two distinct states represented by clusters 10 and 13, as demonstrated by PAGA connectivity and pseudotime analysis. RNA velocity analysis further revealed an alternative to the main NEtD route, suggesting direct transitions from AR^low^ states to early NEPC cluster 10 that bypass the EMT/stem-like cluster 17. Cluster 10 is marked by high expression and activity of ASCL1, a master regulator of SCLC and NEPC (50,54,94–96). In contrast, cluster 13 exhibits neural/neural crest regulons similar to EMT/stem-like cells, along with elevated Notch pathway activity. This is intriguing because Notch signaling promotes self-renewal in neural (97) and NE progenitor cells (98–100) while suppressing ASCL1 expression. Although Notch activation can support NE cell growth (101,102), its role in fate determination between ASCL1+ and ASCL1− NEPC states remains unclear and warrants further investigation. All cellular trajectories ultimately converge at NEPC cluster 7, which lacks ASCL1 and NEUROD1(53) expression, two transcription factors defining major NEPC subtypes. This absence suggests that once NEPC is established, ASCL1 may no longer be required to maintain the neuroendocrine identity. This aligns with recent findings in *Rb1*−/−; *Trp53*−/−; *cMycT58A* organoid-derived tumors, where NEPC persisted without ASCL1 expression(95). Identifying transcriptional drivers that sustain NEPC identity independently of ASCL1 represents a critical area for future research. Furthermore, the detection of NEPC-related GEP1 in bulk RNA-seq patient data and its co-occurrence with other NEPC-associated programs (**Supplementary Fig. 13B**) supports the notion that NEPC represents a continuum of phenotypes rather than discrete subtypes. This aligns with studies showing gradual transcriptional reprogramming during NEtD progression.

### Proliferation Dynamics in NEtD

The identification of proliferative clusters 8 (PRAD) and 3 (NEPC) that maintain strong connectivity to multiple transcriptional states suggests these populations act as reservoirs sustaining phenotypic plasticity and seeding diverse tumor cell lineages. This proliferative flexibility, combined with the existence of alternative paths to NEPC, implies that therapeutic strategies targeting multiple states simultaneously may prove more effective than state-specific interventions. Our findings support the hypothesis that NEtD hijacks embryonic developmental programs to establish therapy-resistant lineages. This is evidenced by the expression of lineage-specific transcription factors across the phenotypic continuum: KLF5 and SOX9 dominate early intermediate states, while MSX1, RUNX3, and TCF7L1 characterize the EMT/stem-like cluster. Early NEPC clusters further activate distinct regulons, including ASCL1, PAX7, and RXRG, which align with known drivers of neuroendocrine plasticity. In bulk RNA-seq analyses, the proliferative signature GEP5 derived from these clusters was detected across all patient subgroups, regardless of AR/NE status (**Supplementary Fig. 15A**). This universal proliferative activity, even in AR-indifferent tumors, highlights a shared vulnerability that could be exploited therapeutically, particularly in the context of epigenetic reprogramming observed during NEtD progression.

### Clinical and Therapeutic Implications

The validation of our findings across multiple independent datasets highlights the robustness and clinical relevance of the LTL331 model for studying NEtD. Using two independent patient-derived scRNA-seq datasets, we identified cellular populations that closely resembled those in LTL331, including PRAD, EMT stem-like and NEPC states. While patient scRNA-seq datasets provide only static snapshots of NEtD, the LTL331 model uniquely captures the temporal progression of cellular states, from late-stage adenocarcinoma (e.g., clusters 4 and 6) and EMT stem-like intermediates to early and terminal NEPC, thereby revealing the dynamic transdifferentiation trajectory that remains unresolved in the patient samples. For example, PRAD cluster 4, which strongly associated with GEP3 and enriched for genes linked to EMT stem-like features and high NEtD potential, identified a subset of CRPC that may reflect ongoing or partial transdifferentiation. This suggests its utility as a prognostic gene signature and a source of potential therapeutic targets. Surprisingly, a motile ciliogenesis transcriptional signature (cluster 6/GEP7) emerges in pre-/early stages and is also detected in both dormant PDX models and androgen-indifferent patient tumors, revealing a previously unrecognized link between cilia biology and prostate cancer progression. Collectively, these findings position LTL331 as a powerful model for tracing clinically meaningful cellular trajectories during NEtD and reveal novel biological programs with prognostic and therapeutic potential.

## Data Availability

The raw data of scRNA-seq for the LTL331 model is currently under review in the SRA and will be updated upon approval. We used publicly available bulk and scRNA-seq data for this study. The processed bulk RNA-seq patient data and metadata are accessible via European Bioinformatics Institute (EMBI-EBI) under accession code E-MTAB-9930 (35). Two scRNA-seq patient datasets include the Gao dataset, available through the NCBI Gene Expression Omnibus Database (GEO) under accession code GSE137829 (11), and the Li dataset accessible from the Genome Sequence Archive for Human (GSA-Human) under accession code HRA002145 (12).

**Table 1.** Summary of seven GEPs and their links to 18 LTL331 clusters and stages.

| GEPs | Stage | Links to 18 clusters |
| --- | --- | --- |
| GEP1 | NEPC | ASCL1 negative: 7, 9, 1, 13, 14, 3 |
| GEP2 | PRAD | 0, 16 |
| GEP3 | PRAD | 4, 5 |
| GEP4 | PRAD | AR<sup>high</sup> clusters from preCX tumor: 11, 12, 15 |
| GEP5 | Proliferative | Cells with high proliferative genes: 3, 8 |
| GEP6 | NEPC | ASCL1 positive: 10 |
| GEP7 | PRAD | Ciliogenesis: 6 |

## Supplementary Data

Supplementary data are available at NAR online.

## Acknowledgements

The graphical abstract includes icons adapted from Bioicons (https://bioicons.com), licensed under CC BY 4.0.

## Author Contributions

**F.S.** performed sequencing, data analysis, and contributed to manuscript writing. **H.C.C.** contributed to data analysis and figures, and writing. **Y.Y.-L.** conducted computational analysis and participated in writing. **D.L.** was responsible for PDX model development and pathology analysis. **T.M., S.V., D.O.** assisted with data analysis. **A.H.** performed sequencing. **R.B.** provided expertise in biostatistics. **S.L.** managed the project. **H.X.**, **R.W.**, and **X.D.** supported PDX model work. **J.L.** analyzed and generated the processed datasets. **H.J.-K.** contributed to scientific discussion and manuscript writing. **M.E.G.** provided patient tumor samples and clinical data. **N.A.L.** contributed to data analysis and manuscript writing. **Y.W.** led PDX model development and conceptualization. **C.C.C.** led the genomics analysis, conceptualized the study, and contributed to writing. **Y.W.** and **C.C.C.** jointly supervised the project and mentored trainees. All authors made intellectual contributions.

## Funding

This research was partially supported by the Canadian Institutes of Health Research (CIHR) under grants #153081, #173338, #180554, and #186331 awarded to Y.W.; the Terry Fox Research Institute (grant #1109) awarded to C.C.C. and Y.W.; and a Canadian Cancer Society Breakthrough Team Grant generously supported by the Lotte & John Hecht Memorial Foundation (CCS grant #707683) awarded to Y.W. Additional support was provided by a BC Cancer Foundation grant (1PRRG012) to Y.W. C.C.C. acknowledge support from CIHR grants PJT-175238 and PJT-153073. H.J.K. acknowledges funding from the Ministry of Health and Welfare (MOHW 114-TDU-B222-134002) and the National Science and Technology Council (NSTC-114-2634-039-001 and 114-2326-B-038-001). N.A.L. was supported by funding from TUBITAK (221Z116), the U.S. Department of Defense (W81XWH-21-1-0234), and CIHR (PJT-173331).

## Notes

### Competing Interest Statement

The authors have declared no competing interest.

### Summary of Updates

Adding new authors included in the manuscript but missing in the metadata

