## Supplementary material for "Longitudinal single-cell RNA sequencing of a neuroendocrine transdifferentiation model reveals transcriptional reprogramming in treatment-induced neuroendocrine prostate cancer": Supplementary Figures.pdf

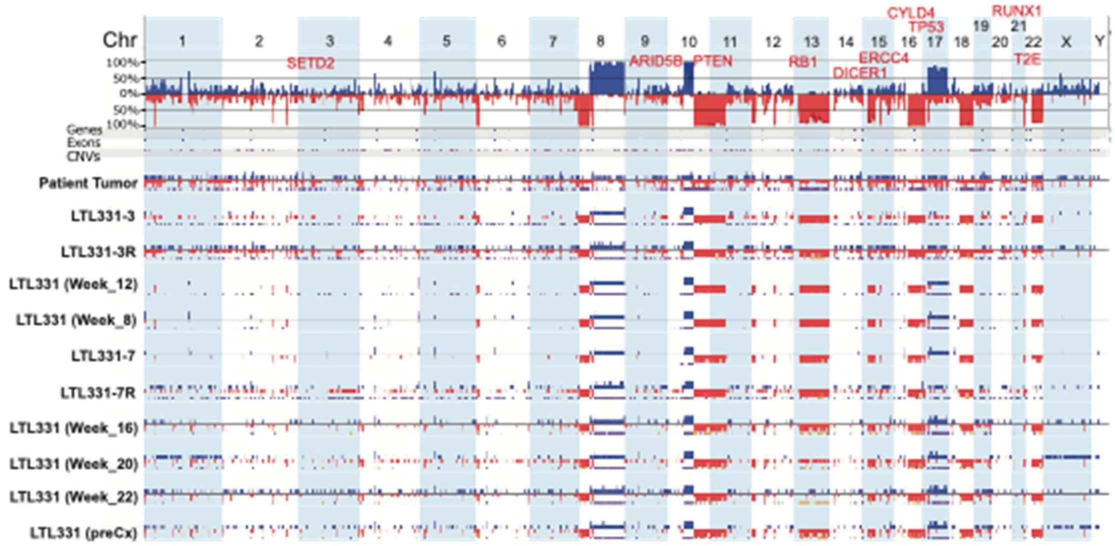


***Supplementary Figure 1.*** Copy number profile of the parental patient tumor (927) and LTL331 model. Passages including parentheses behind are the ones used for scRNA-seq studies, and passages 3 and 7 neuroendocrine samples are marked with a letter "R" (LTL331-3R, LTL331-7R). The graph with percentages in the top is the frequency plot, which shows how often copy number losses (red) and gains (blue) are observed in a given region in all samples, and the height of the color bar is proportional to the percentage of samples where a given aberration was observed. The frequency for each sample is presented below, color-coded in the same way. The height of a bar is proportional to the scale of a rearrangement – a single copy gain/heterozygous loss is half-height, high-copy gain/homozygous loss is presented by the full-length bar. Colored bands below each graph demarcate regions of loss of heterozygosity (orange-brown; e.g., Chr 13 in LTL331 Week 22) and allelic imbalance (purple; e.g., Chr 8 across all samples) detected by Nexus algorithms.

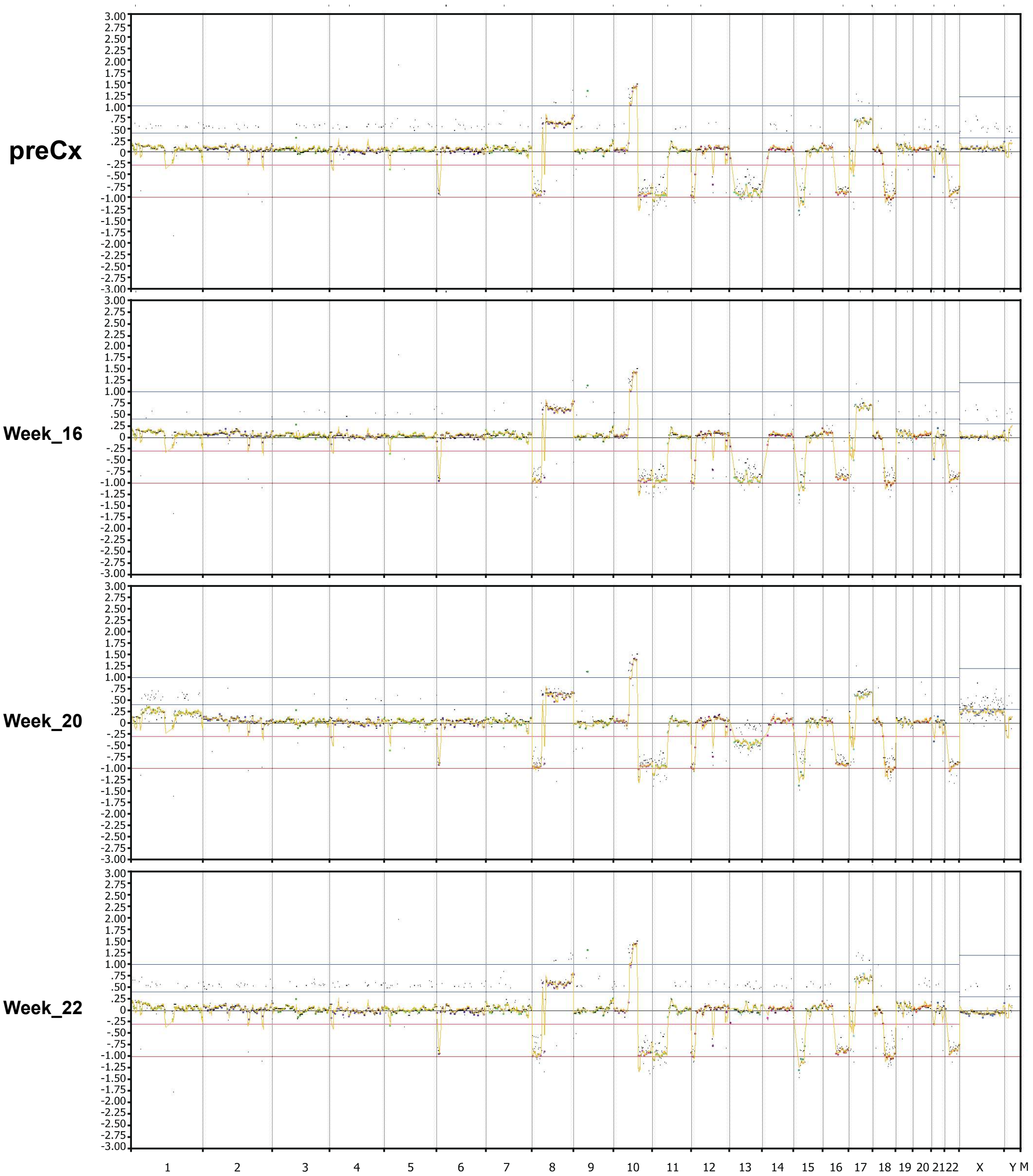


***Supplementary Figure 2.*** WES-based copy number profiles for a subset of L TL331 model (preCx, week 16, 20, 22). The orang line represents smoothed CN profile inferred from WES data. Chromosomes are marked across the bottom, and chromosome order is as expected. The inferred “normal” copy number is represented by the horizontal blue lines, therefore the orange line above the blue line demarcates regions of copy number gains, and below the regions of copy number losses. All four plots are overall concordant, e.g. the characteristics such as Chr 8q gain, 16q loss, etc, can be detected in all samples.

A

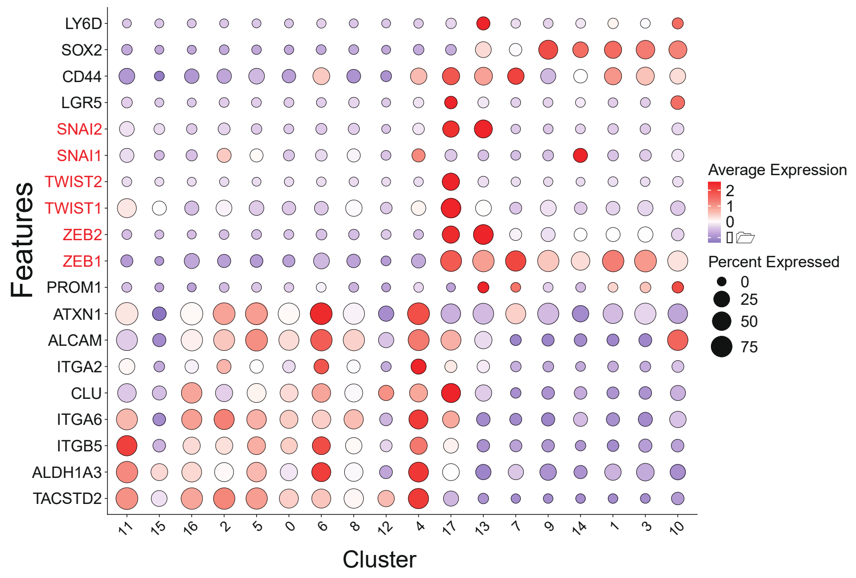


B

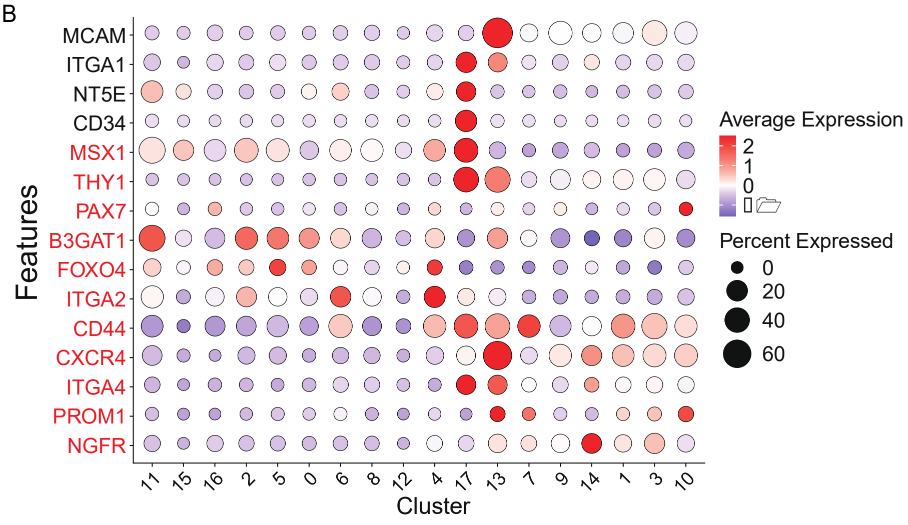


***Supplementary Figure 3.*** Expression of EMT, stem cell and progenitor related genes in L TL331. (A) Dot plot displaying the expression of stem ce ll and progenitor related gene sets, and master regulators of EMT (written in red) (B) mesenchym al and neural crest stem cell genes (written in red) for each cluster. The size of the dot represents the percentage of cells in the cluster express - ing the gene and color shows the level of expres sion.

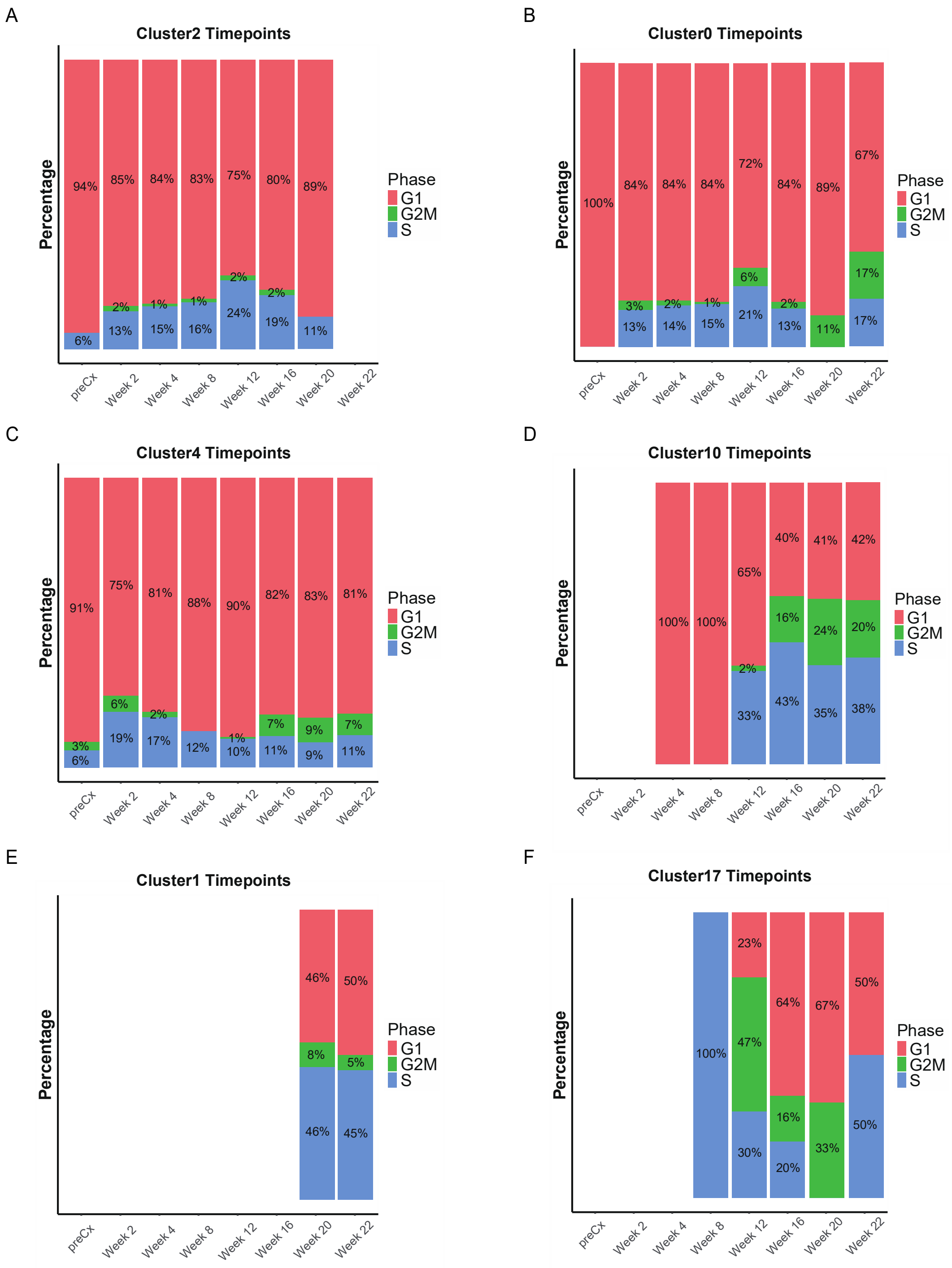


***Supplementary Figure 4.*** Cell cycle profiles of selected clusters across time-points. (A)-(F). Percentage of cells in G1, G and G2/M phases of the cell-cycle for selected clusters at each timepoint during NEtD.

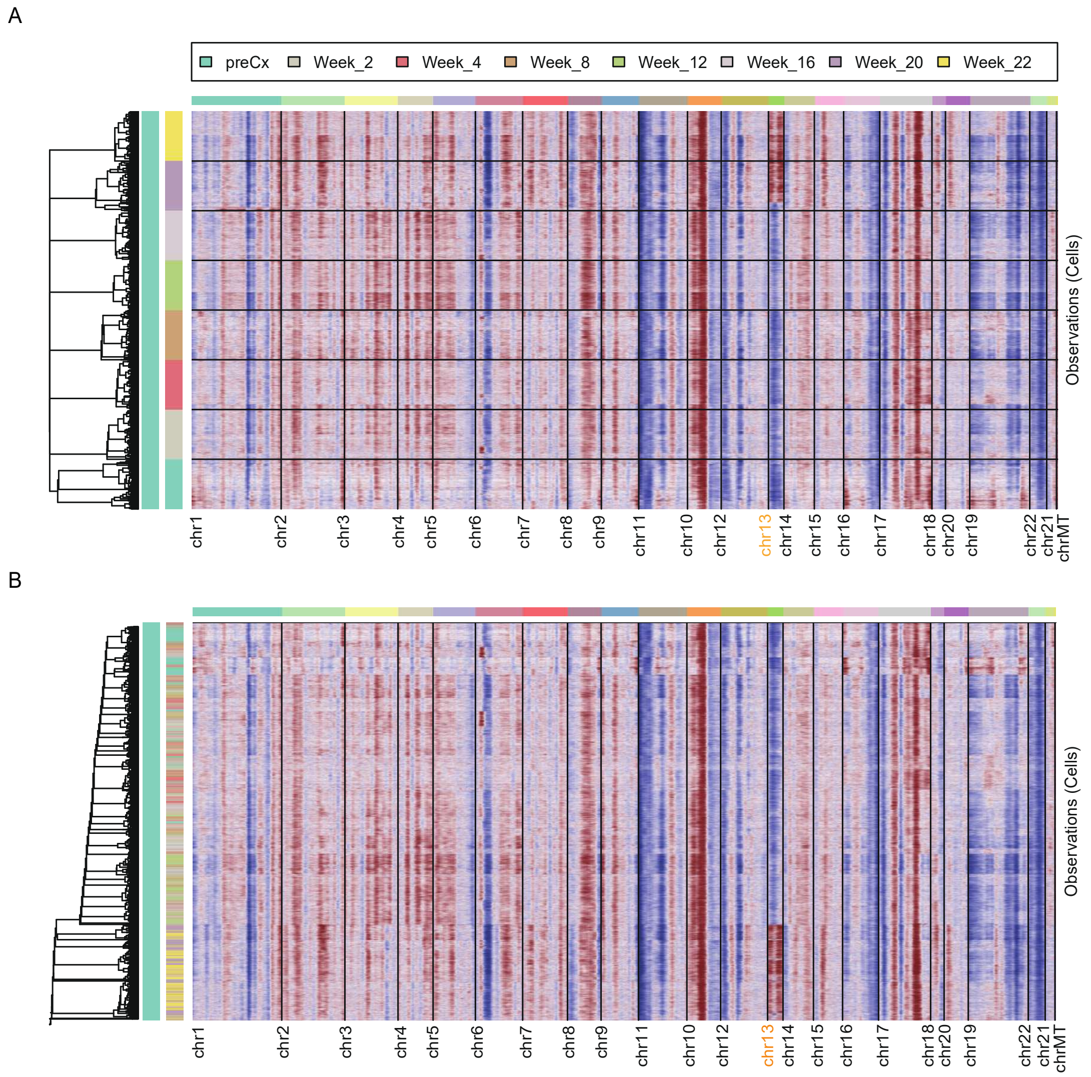


***Supplementary Figure 5.*** InferCNV profile of the LTL331 model grouped by time-points. (A) InferCNV profiles of the cells are grouped by t he time-point at which they are were collect - ed. (B) InferCNV profiles of the cells without grou ping by time-point. Cells are colored based on the time points at which they were collected, with the corresponding color scheme displayed above the graph. Chromosomes are indicated at the bottom of the plots in black. The inferred copy number losses are shown in blue, while gain s are shown in red. Each row represents one cell, in panel A, the cell rows are clustered first by sample and subsequently by similarity of their CNV profiles within each sample, in panel B , the cells are clustered based on their similar - ity in CNV profiles. The dendrogram on the left di splays the clustering results.

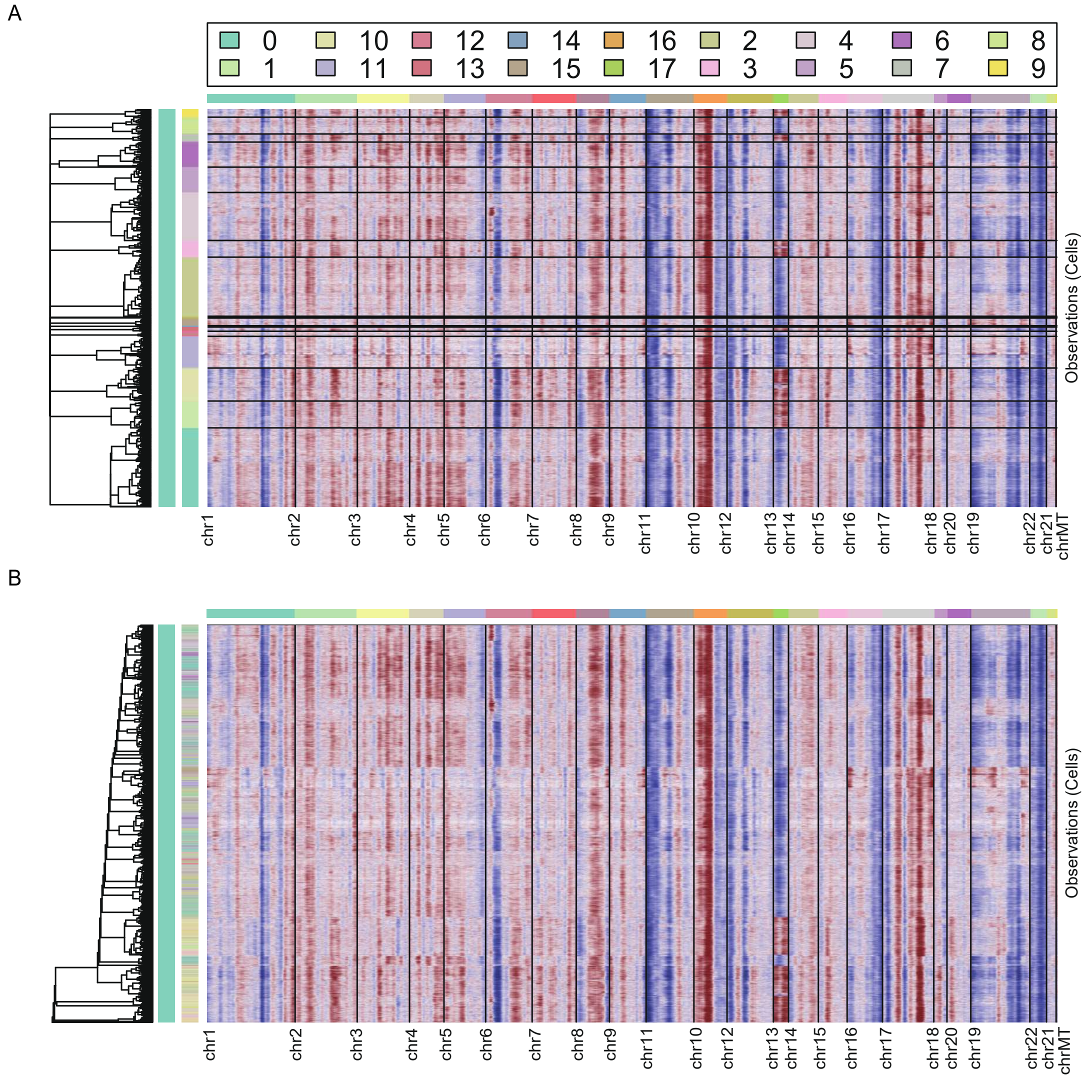


***Supplementary Figure 6.*** InferCNV profile of the LTL331 clusters.
(A) InferCNV profiles of clusters. (B) InferCNV profiles of the cells independent of the clusters that they are derived from. Cells are colored base d on the time points at which they were collected, with the corresponding color scheme d isplayed above the graph. Chromosomes are indicated at the bottom of the plots in black. The inferred copy number losses are shown in blue, while gains are shown in red. Each row repr esents one cell, in panel A, the cell rows are clustered first by original cluster assignment and subsequently by similarity of their CNV profiles within each cluster, in panel B, the cells are clustered based on their similarity in CNV profiles independent of their original cluster assig nments. The dendrogram on the left displays the clustering results.

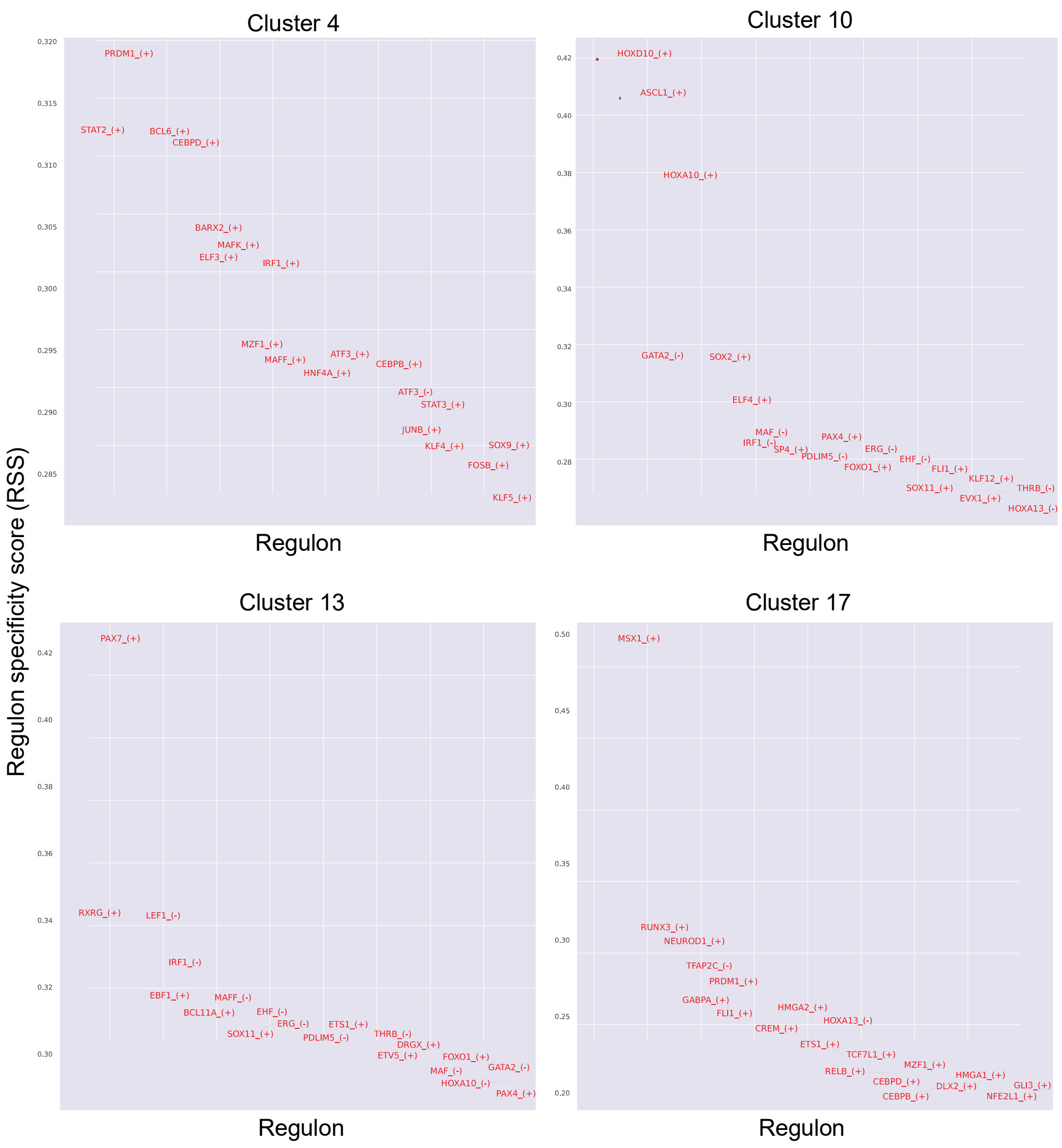


***Supplementary Figure 7.*** Regulon specificity scores (RSS) of selected clusters.
Regulon specificity scores (RSS) of the top 20 regulons that are unique to clusters 4, 10, 13 or 17.

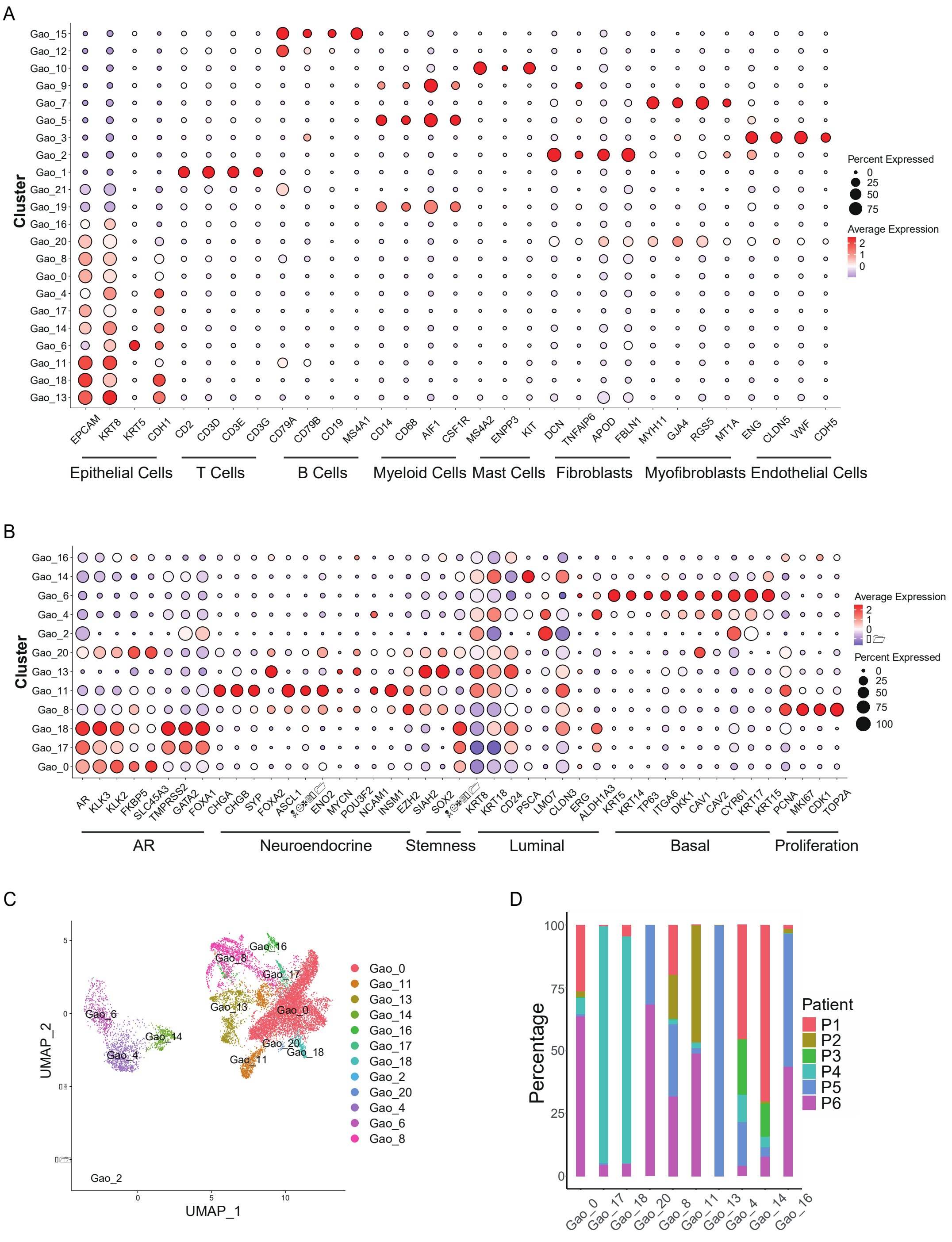

***Supplementary Figure 8.*** Analysis of the first patient-derived scRNA-seq dataset (Gao dataset). (A) Dot plot showing the expression prevalence of canonical markers across all clusters detected in the first public scRNA-seq dataset 20. (B) Dot plot displaying the expression prevalence of canonical lineage signature marker genes across epithelial clusters. (C) UMAP visualization of epithelial cells extracted for further analysis. (D) Proportions of patient-derived cells contributing to each epithelial cluster in this dataset.

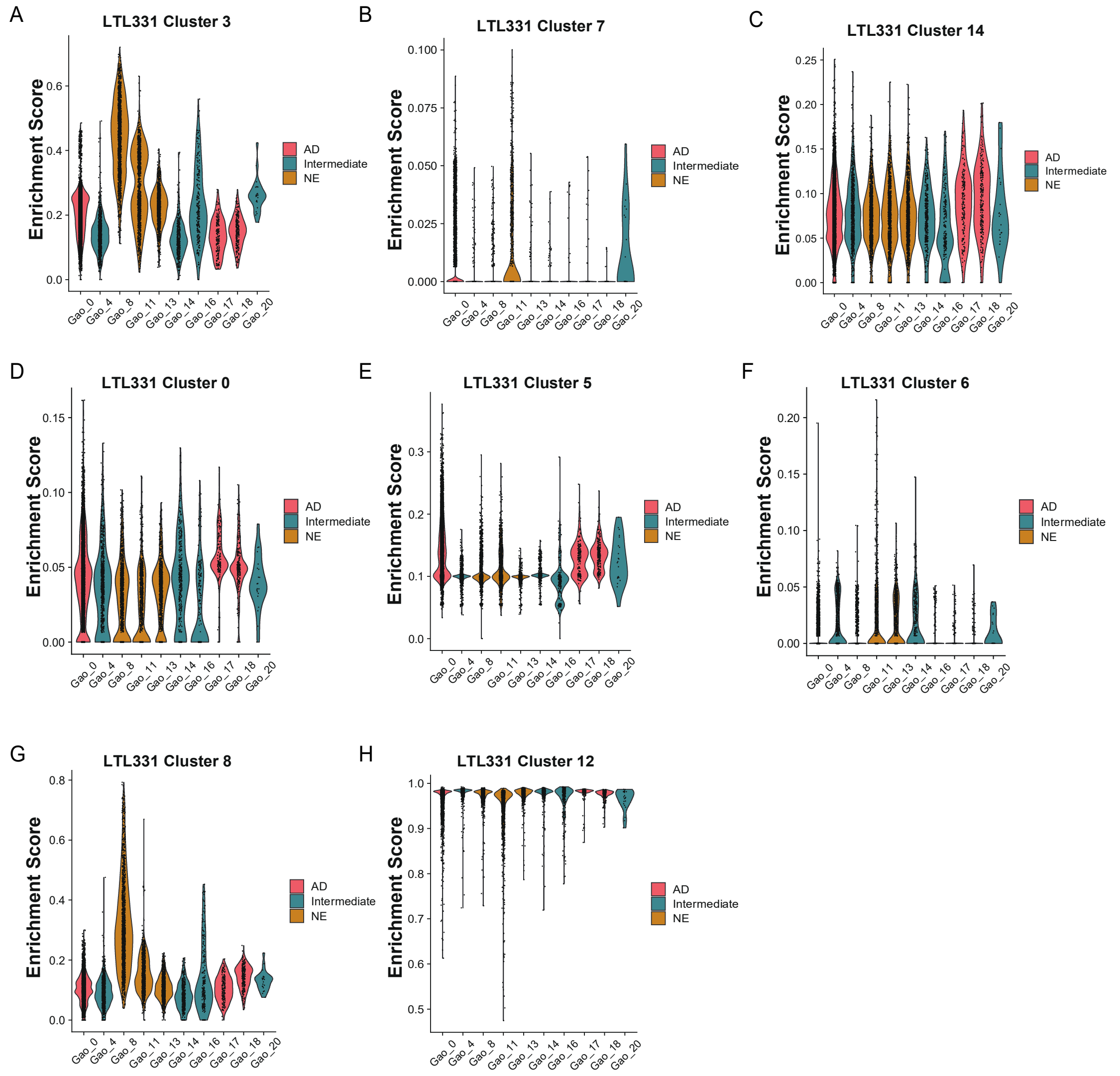


***Supplementary Figure 9.*** Scoring of PRAD and NEPC LTL331 cluster-specific genes in the first patient-derived scRNA-seq dataset (Gao dataset).
(A)-(H) Violin plots displaying the enrichment scores for PRAD and NEPC L TL331 cluster-specific gene signatures across the first public scRNA-seq dataset 20 that were not included in Fig. 6.

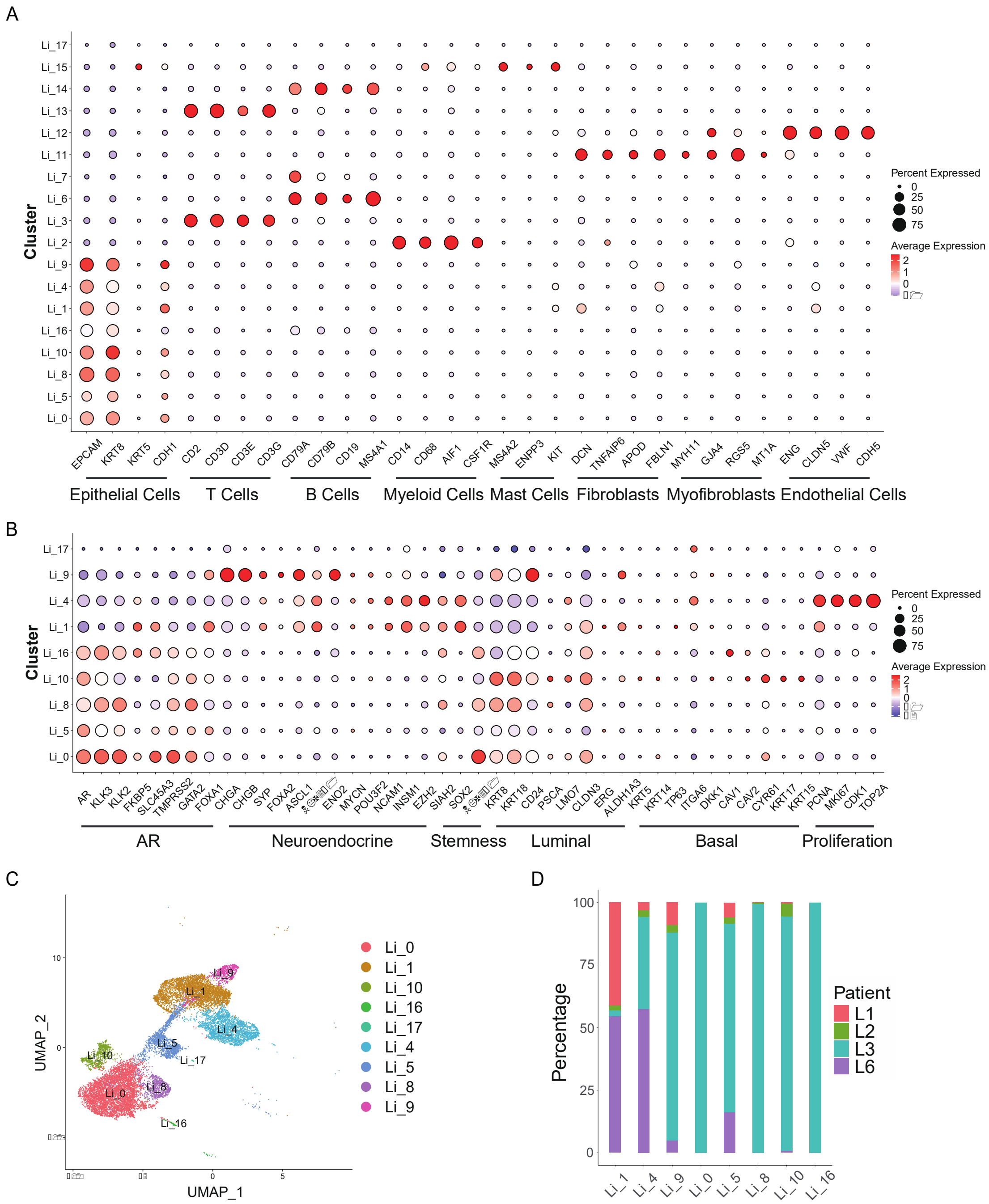


***Supplementary Figure 10.*** Analysis of the second patient-derived scRNA-seq dataset (Li dataset). (A) Dot plot displaying the expression prevalence of canonical markers across all clusters detected in the second public scRNA-seq dataset 21 (W ang, Wang et al. 2022). (B) Dot plot showing the expres - sion prevalence of canonical lineage signature marker genes across epithelial clusters in this dataset. (C) UMAP visualization of epithelial cells extracted for further analysis. (D) Proportions of patient derived cells contributing to each epithelial cluster in this dataset.

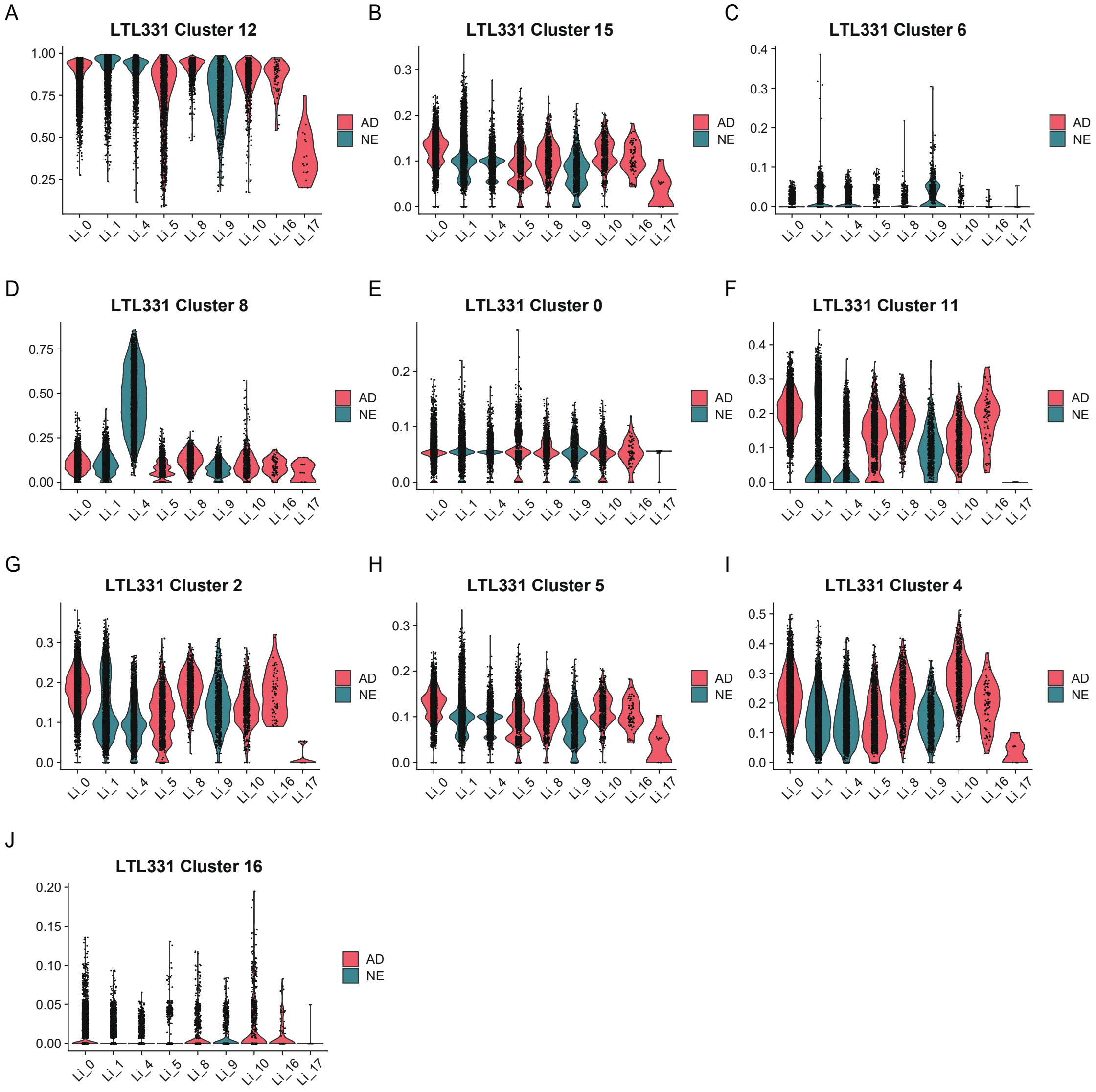


***Supplementary Figure 11.*** Scoring of LTL331 PRAD cluster-specific genes in the second patient-de - rived dataset (Li dataset).
(A)-(J) Violin plots illustrating enrichment scores for PRAD L TL331 cluster-specific gene signatures across the second public scRNA-seq dataset 21.

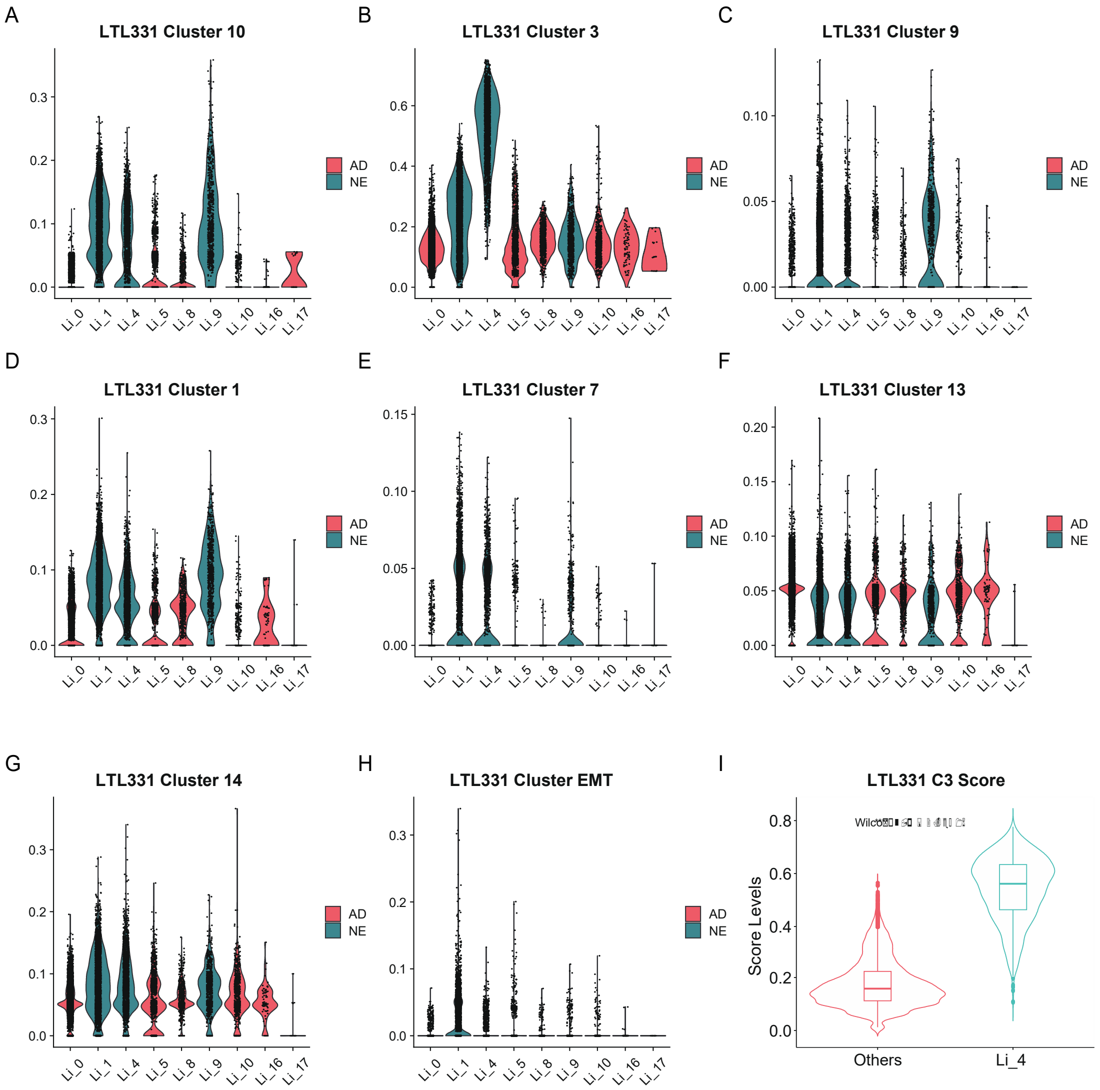


***Supplementary Figure 12.*** Scoring of LTL331 NEPC and EMT cluster-specific genes in the second patient-derived dataset (Li dataset).

(A)-(H) Violin plots showing enrichment scores for NEPC/EMT LTL331 cluster-specific gene signatures across the second public scRNA-seq dataset 21. (I) V iolin plot depicting enrichment scores of the LTL331cluster3 specific gene signature in cluster 4 compared to all other clusters within the dataset. The significance is determined by the Wilcoxon test, the boxplot represents the interquartile range (IQR)

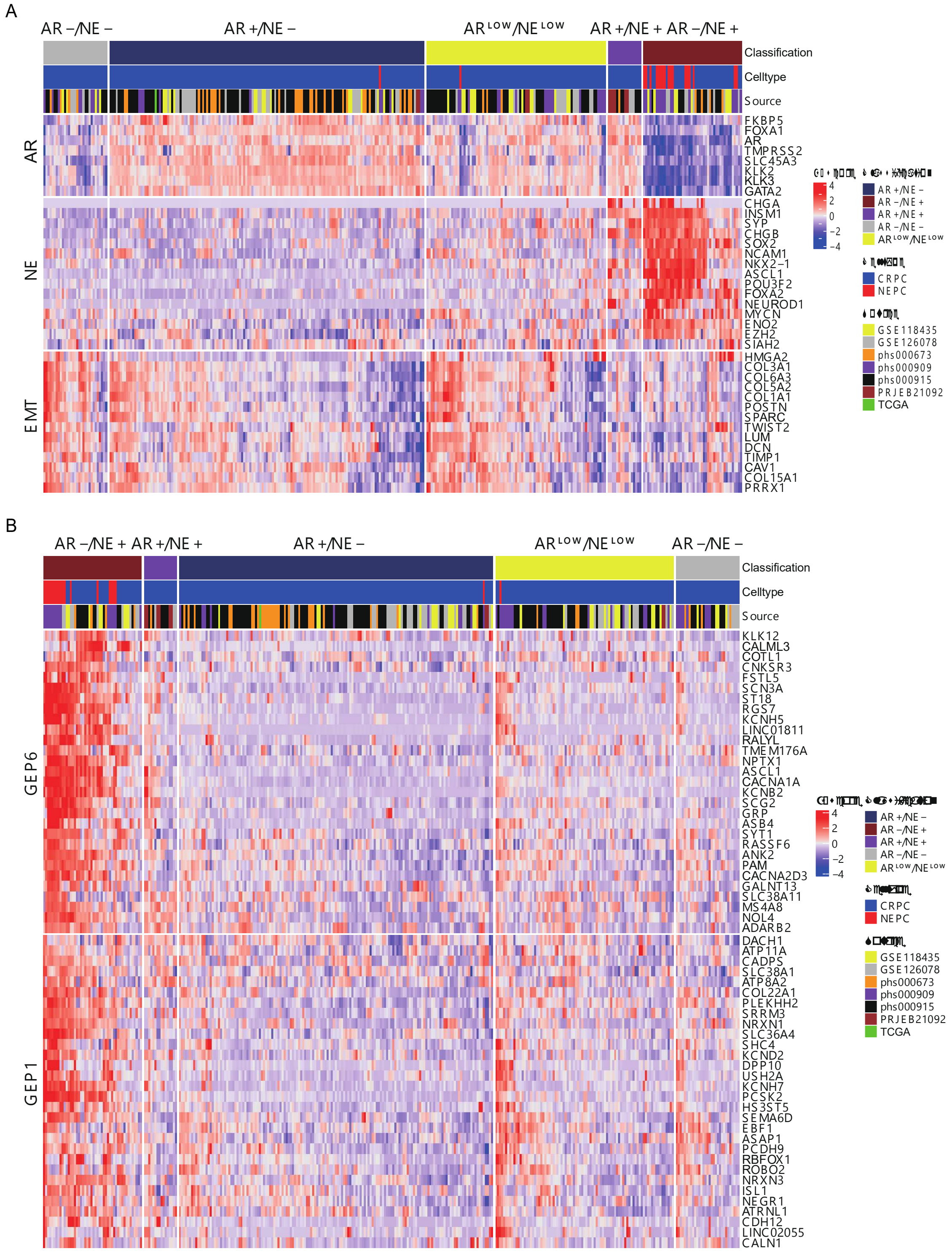
A
AR −/NE −
AR +/NE −
AR LOW/NE LOW
AR +/NE +
AR −/NE +
Classification
Celltype
Source
AR
NE
EMT
FKBP5
FOXA1
AR
TMPRSS2
SLC45A3
KLK2
KLK3
GATA2
CHGA
INSM1
SYP
CHGB
SOX2
NCAM1
NKX2−1
ASCL1
POU3F2
FOXA2
NEUROD1
MYCN
ENO2
EZH2
SIAH2
HMGA2
COL3A1
COL6A3
COL5A2
COL1A1
POSTN
SPARC
TWIST2
LUM
DCN
TIMP1
CAV1
COL15A1
PRRX1
4
2
0
−2
−4
AR +/NE −
AR −/NE +
AR +/NE +
AR −/NE −
AR LOW/NE LOW
CRPC
NEPC
GSE118435
GSE126078
phs000673
phs000909
phs000915
PRJEB21092
TCGA
B
AR −/NE +
AR +/NE +
AR +/NE −
AR LOW/NE LOW
AR −/NE −
Classification
Celltype
Source
GEP6
GEP1
KLK12
CALML3
COTL1
CNKSR3
FSTL5
SCN3A
ST18
RGS7
KCNH5
LINC01811
RALYL
TMEM176A
NPTX1
ASCL1
CACNA1A
KCNB2
SCG2
GRP
ASB4
SYT1
RASSF6
ANK2
PAM
CACNA2D3
GALNT13
SLC38A11
MS4A8
NOL4
ADARB2
DACH1
ATP11A
CADPS
SLC38A1
ATP8A2
COL22A1
PLEKHH2
SRRM3
NRXN1
SLC36A4
SHC4
KCND2
DPP10
USH2A
KCNH7
PCSK2
HS3ST5
SEMA6D
EBF1
ASAP1
PCDH9
RBFOX1
ROBO2
NRXN3
ISL1
NEGR1
ATRNL1
CDH12
LINC02055
CALN1
4
2
0
−2
−4
AR +/NE −
AR −/NE +
AR +/NE +
AR −/NE −
AR LOW/NE LOW
CRPC
NEPC
GSE118435
GSE126078
phs000673
phs000909
phs000915
PRJEB21092
TCGA

***Supplementary Figure 13.*** Reclassification of public Bulk RNA-seq PCa patient samples and NEPC GEP gene expression.
(A) PCa patients' samples were reclassified as AR+/NE-, AR-low/NE-low, AR+/NE+ and AR-/NE+ subgroups according to hierarchal clustering of AR pathway and NE marker gene expression. The bottom panel displays the expression of the top 15 genes differentially expressed in EMT stem-like cluster 17. (B) Heatmaps showing the expression patterns of the top30 genes from GEP1 (ASCL1-) and GEP6 (ASCL1+) genes across subtypes. Original patient classification and the original sources were color-coded at the top and annotated on the right. Gene expression values are presented as z-scaled,.

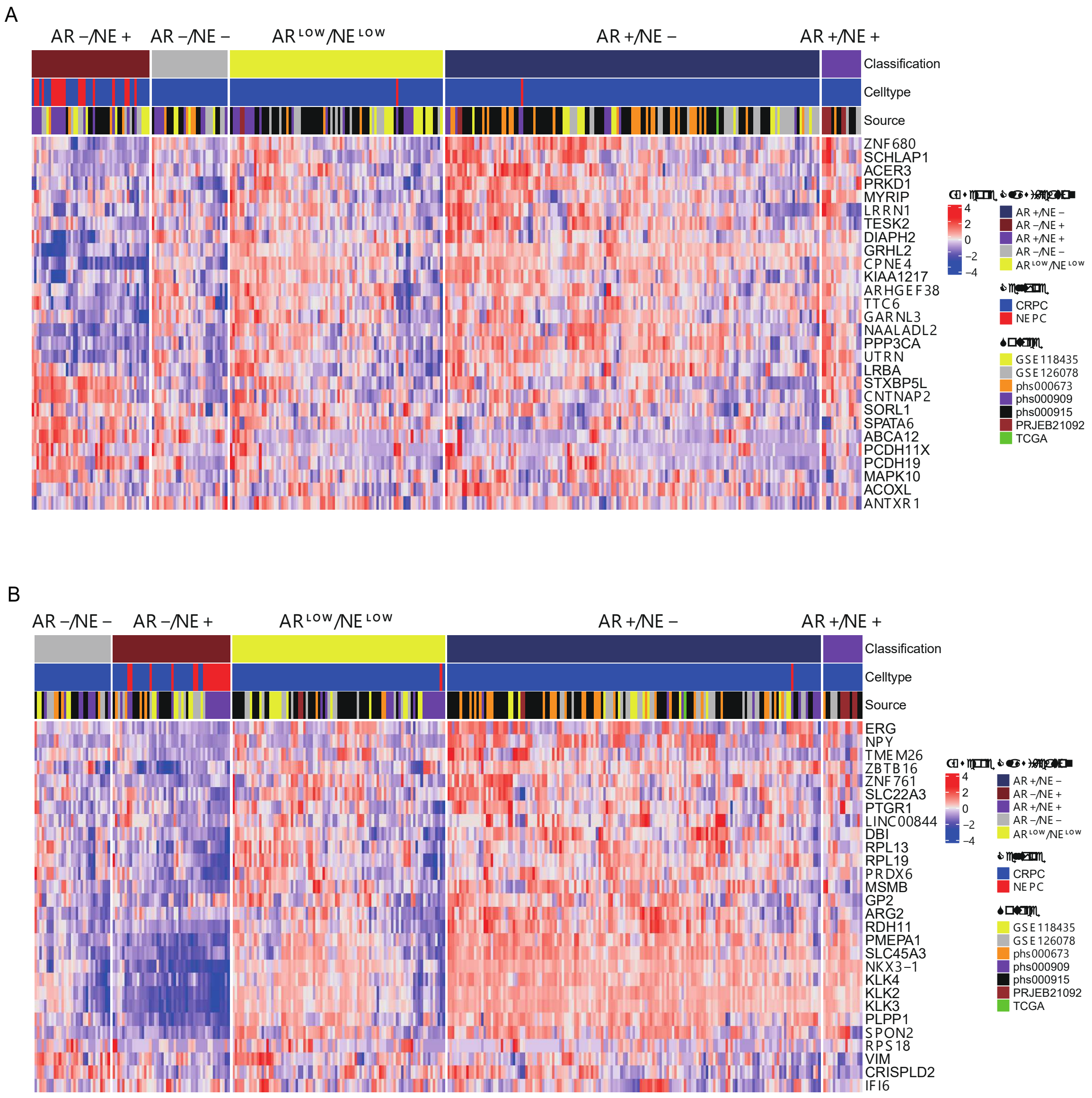


***Supplementary Figure 14.*** Expression of GEP2 and 4 across PCa patients.
Heatmaps showing the expression of the top 30 genes from GEP2 (A) and GEP4 genes (B) across PCa subgroups. The original patient classification and the original sources were color-coded at the top and annotated on the right. Gene expression values are presented as z-scaled.

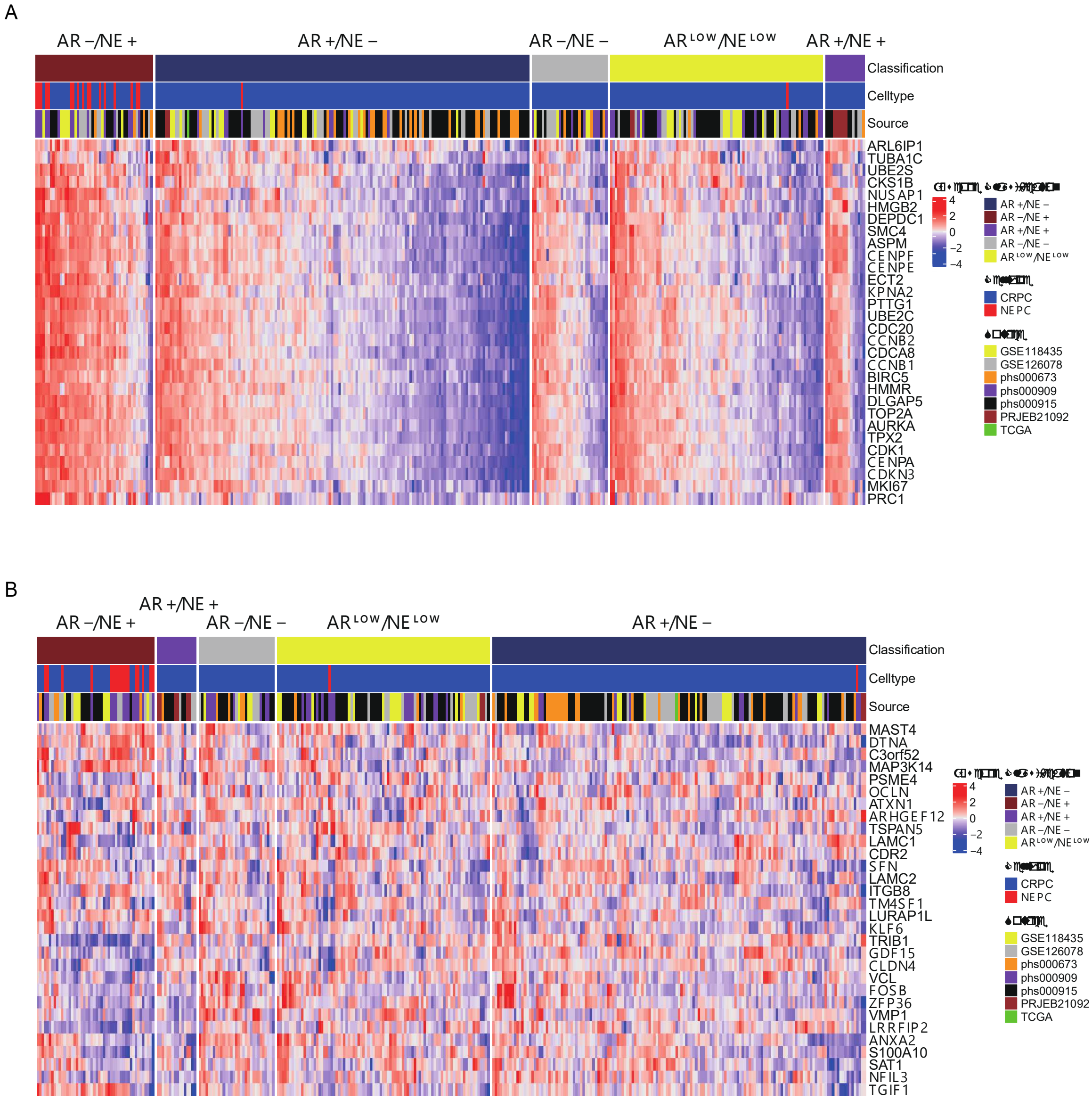


***Supplementary Figure 15.*** Expression of GEP5 and 3 across PCa patients.
Heatmaps showing the expression of the top 30 genes from GEP5 (A) and GEP3 genes (B) across PCa subgroups. The original patient classification and the sources were color-coded at the top and annotated on the right. Gene expression values are presented as z-scaled.

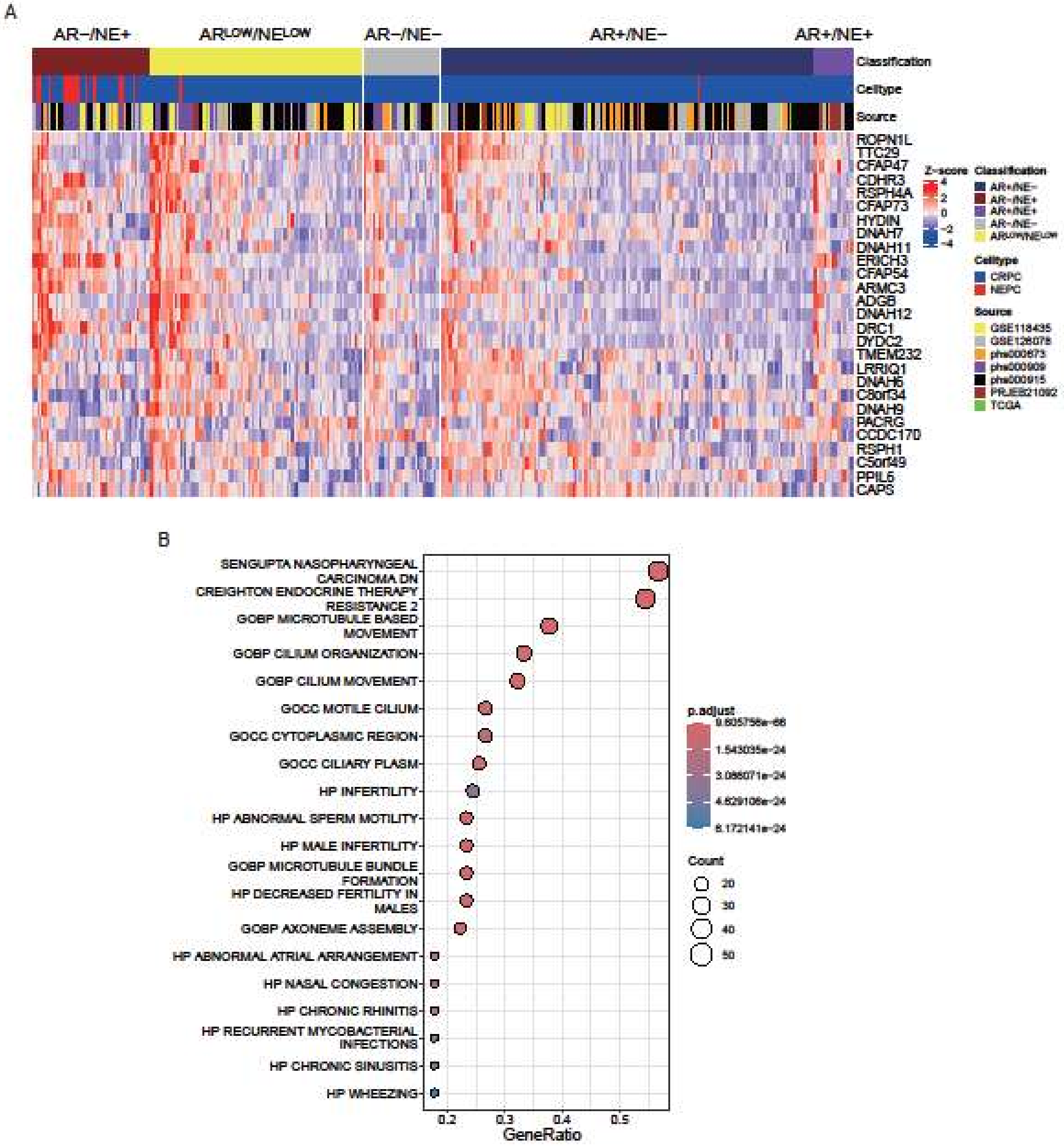


***Supplementary Figure 16.*** Analysis of genes from GEP7.

(A) Heatmap depicting the expression of the top 30 genes from GEP7. The original patient classification and the sources were color-coded at the top and annotated on the right. Gene expression values are presented as z-scaled, (B) Over-representation analysis of all genes from GEP7 genes using Hallmark, GO, KEGG, and Reactome gene sets.

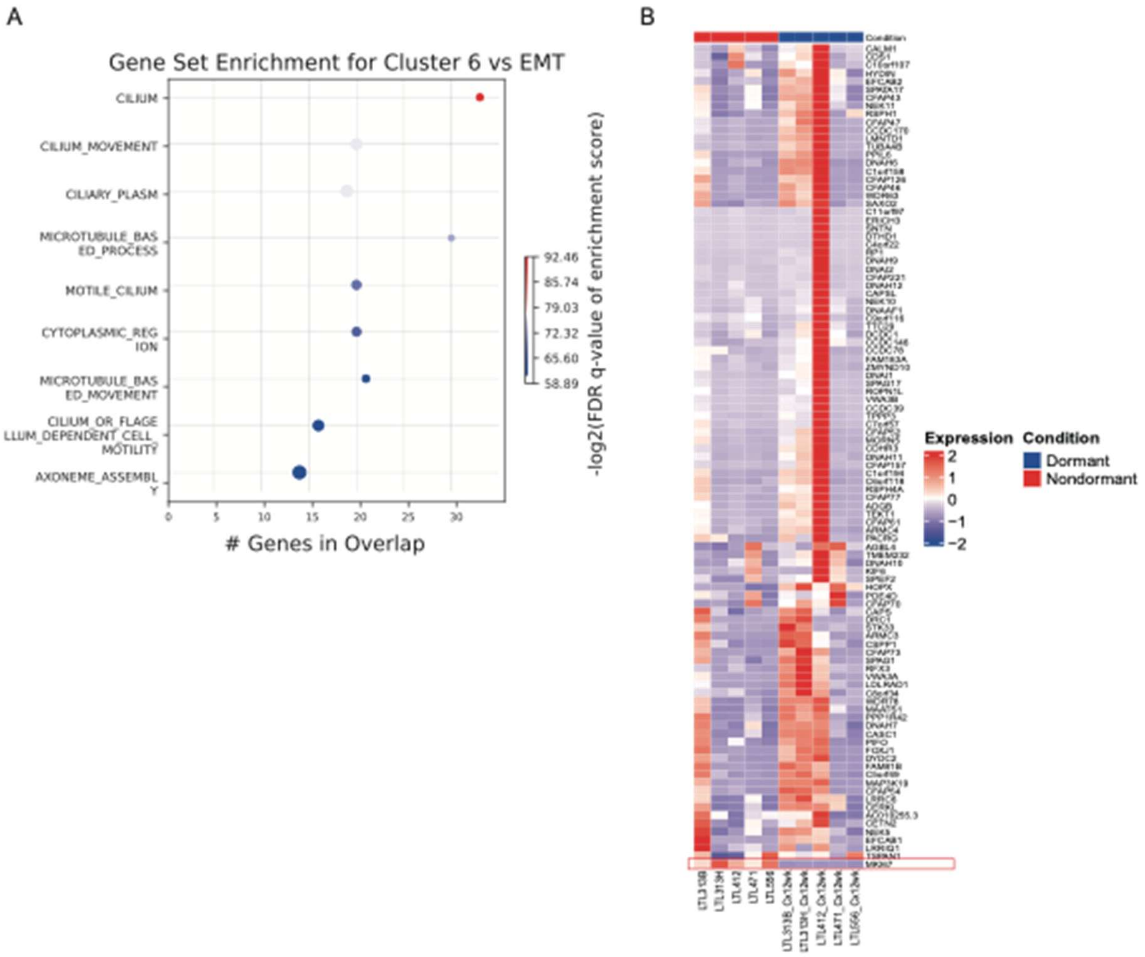


***Supplementary Figure 17.*** GSEA analysis of cluster 6 and validation of GEP7 in dormant PDX models.
(A) Gene set enrichment analysis of differentially expressed genes in cluster 6 compared to cluster 17 and (B) Heatmap showing expression of the 100 GEP7 genes and proliferation marker MKI67 (bottom) across five published dormant (blue) and non-dormant (red) PDX models.
